# Heparan sulfate modifications of betaglycan promote TIMP3-dependent ectodomain shedding to fine-tune TGF-β signaling

**DOI:** 10.1101/2023.08.29.555364

**Authors:** Alex S Choi, Laura M Jenkins-Lane, Wade Barton, Asha Kumari, Carly Lancaster, Calen Raulerson, Hao Ji, Diego Altomare, Mark D Starr, Regina Whitaker, Rebecca Phaeton, Rebecca Arend, Michael Shtutman, Andrew B Nixon, Nadine Hempel, Nam Y Lee, Karthikeyan Mythreye

## Abstract

In pathologies such as cancer, aberrant Transforming Growth Factor-β (TGF-β) signaling exerts profound tumor intrinsic and extrinsic consequences. Intense clinical endeavors are underway to target this pivotal pathway. Central to the success of these interventions is pinpointing factors that decisively modulate the TGF-β responses. Betaglycan/type III TGF-β receptor (TβRIII), is an established co-receptor for the TGF-β superfamily known to bind directly to TGF-βs 1-3 and inhibin A/B. While betaglycan can be membrane-bound, it can also undergo ectodomain cleavage to produce soluble-betaglycan that can sequester its ligands. The extracellular domain of betaglycan undergoes heparan sulfate and chondroitin sulfate glycosaminoglycan modifications, transforming betaglycan into a proteoglycan. Here we report the unexpected discovery that the heparan sulfate modifications are critical for the ectodomain shedding of betaglycan. In the absence of such modifications, betaglycan is not shed. Such shedding is indispensable for the ability of betaglycan to suppress TGF-β signaling and the cells’ responses to exogenous TGF-β ligands. Using unbiased transcriptomics, we identified TIMP3 as a key regulator of betaglycan shedding and thereby TGF-β signaling. Our results bear significant clinical relevance as modified betaglycan is present in the ascites of patients with ovarian cancer and can serve as a marker for predicting patient outcomes and TGF-β signaling responses. These studies are the first to demonstrate a unique reliance on the glycosaminoglycan modifications of betaglycan for shedding and influence on TGF-β signaling responses. Dysregulated shedding of TGF-β receptors plays a vital role in determining the response and availability of TGF-βs’, which is crucial for prognostic predictions and understanding of TGF-β signaling dynamics.

## Introduction

Type III TGF-β receptor (TβRIII) / betaglycan (BG) is a widely expressed transmembrane proteoglycan and an established coreceptor for a subset of the TGF-β superfamily of ligands ^1,2^. As a co-receptor, BG can either increase or decrease signaling by TGF-β superfamily members that directly bind BG including all isoforms of TGF-β1,2 and 3 ^3,4^, as well as BMP2, 4, 7 ^5,6^, GDF- 5 ^6^, Inhibin A ^7^, and Inhibin B ^8^. Betaglycan binds TGF-β2 with greater affinity than TGF-β1 or TGF-β3 ^9^ and thus cells lacking BG expression do not respond as well to TGF-β2 as compared to TGF-β1 and TGF-β3 in equimolar settings, requiring up to 500-fold higher concentrations of TGF-β2 to achieve the same potency of activation as TGF-β1 and TGF-β3 in the absence of BG ^10–12^. These observations are not cell-line specific and have been reported in multiple cell types including in cancer cell lines and non oncogenic models ^13–15^. BG likely functions to bind TGF-β2 and concentrate the ligand to facilitate access to the Type II-TGF-β receptor kinase, effectively restoring cellular sensitivity to TGF-β2 to comparable levels of TGF-β1/3 ^3^. Both BG and TGF-β2 knockout mice show similar defects in vivo ^16,17^, demonstrating the physiological reliance of TGF- β2 on BG. In the case of TGF-β1 responsiveness, BG can either stimulate or inhibit TGF-β1 signaling ^18–21^.

Transmembrane proteoglycans including betaglycan can be proteolytically cleaved, releasing soluble ectodomain into the ECM in a process called ectodomain shedding ^22^. Only 2 to 4% of cell surface molecules undergo shedding ^23,24^, and dysregulated shedding is associated with various pathologies including cancer ^22,24,25^ suggesting that maintaining the levels of shed proteins may be critical. Notably, in women’s cancers including ovarian ^26^, breast ^19^, as well as granulosa cell tumors that arise from ovarian sex cord-stromal cells^27^, lower BG expression in the tumor cells as compared to adjacent normal tissue has been reported ^27–29^ with lower BG expression found to be an indicator of poor patient outcomes ^26,30^. Previous studies also indicated that shed-BG in tumor cells reduces their cell migration, invasion, and metastasis properties ^19,21,31^. Similar observations have been made in normal epithelial cells as well ^32^. Compared to membrane-bound BG, shed-BG can reduce TGF-β signaling ^33,34^. General regulators of proteoglycan shedding include proteases (also called sheddases) of the ADAM/ADAM-related family of proteins ^35,36^ and MMPs, most notably, but not limited to MMPs-1,2, and 7 ^35,37–40^, as well as chemical agents such as phorbol ester and calcium ionophores ^36,38,39^, and specific serum factors ^22,24,25,38,41^. BG shedding is however unaffected by phorbol esters, calcium ionophores, PMA, and serum factors ^23,42^, but is stimulated by pervanadate ^38^, and is inhibited by TAPI-2, an MT-MMP/ADAM protease inhibitor ^43^.

A distinctive feature of BG lies in the extracellular domain modifications with glycosaminoglycan (GAG) chains at serine residues Ser^534^ and Ser^545^, to which heparan sulfate (HS) and chondroitin sulfate (CS) chains are covalently attached ^44–49^. HS is a repeating unit of N-acetylglucosamine and glucuronic acid and CS is a repeat of n-acetyl-galactosamine and glucuronic acid ^50^. BG is commonly referred to as a “part-time proteoglycan” since BG can be expressed on the cell surface with or without GAG chains ^51–54^. Previous reports suggest that GAG chains are not essential for the ligands that bind to the core domain of BG ^9,44^, however, GAG chains of BG can mediate the binding of growth factors such as FGF2 ^55^ and Wnt3A ^49^, where Wnt3A signaling can be influenced by the HS and CS chains ^49^. In addition to affinities of ligands to the GAG chains of BG, the presence of the GAG chains has also been proposed to prevent access of the BG core binding ligands to their respective signaling receptors, suggesting disruption of TGF-β signaling and function ^48,56^.

While previous studies have demonstrated the effect of overall BG levels on TGF-β signaling, a thorough understanding of the role of BG GAG chains and their influence on regulating TGF-β signaling if any, is currently lacking. Here, we sought to address this. We report a direct role for the BG GAG modifications on ectodomain shedding of BG and identify BG modification-specific expression changes in TIMP3, a negative regulator of BG ectodomain shedding. We also demonstrate that the presence of GAG modifications on BG is critical for fine-tuning TGF-β signaling and invasive properties of tumor cells. Lastly, we report that higher amounts of shed- BG in the ascites fluid of ovarian cancer patients correlate with advanced-stage disease and serve as a negative predictor of patient survival.

## Results

### TβRIII / betaglycan (BG) glycosaminoglycan modifications promote ectodomain shedding

To address whether glycosaminoglycan modifications on BG impact ectodomain shedding, we used a previously generated double Ser- Ala point mutation at S534 and S545 of BG, to eliminate the GAG chain attachment sites on BG ^44–49^. Expression of this construct has been reported to prevent GAG attachment ^38,44,55,57^ and was recapitulated in cancer cells used here that represent a spectrum of epithelial ovarian cancers (**Suppl. Fig 1A**). HEYA8, SKOV-3, and OVCA-429 ovarian cancer (OVCA) cell lines were chosen due to low endogenous expression of BG (**Suppl. Fig 1B**) and BG constructs (Full-Length (BG-FL) or BG lacking GAG chains (BG-ΔGAG) were either transiently or stably overexpressed (**Fig 1A, Suppl. Fig 1A**). Previous studies have reported no effects of the GAG modifications on BG on its ability to bind TGF-β ^44,48^ and this was confirmed here as well using cell surface receptor [^125^ I]-TGF-β1 binding and crosslinking (**Fig 1A.i**). Cell surface binding of BG to TGF-β confirmed that FL-BG cells present GAG modifications ranging from ∼90kDa to 250kDa (**Fig 1A.i second lanes**), while ΔGAG-BG expressing cells which are competent at binding TGF-β showed BG devoid of high molecular weight GAG chains with a core band at ∼95kDa (**Fig 1A.i third lanes**).

**Fig 1.**
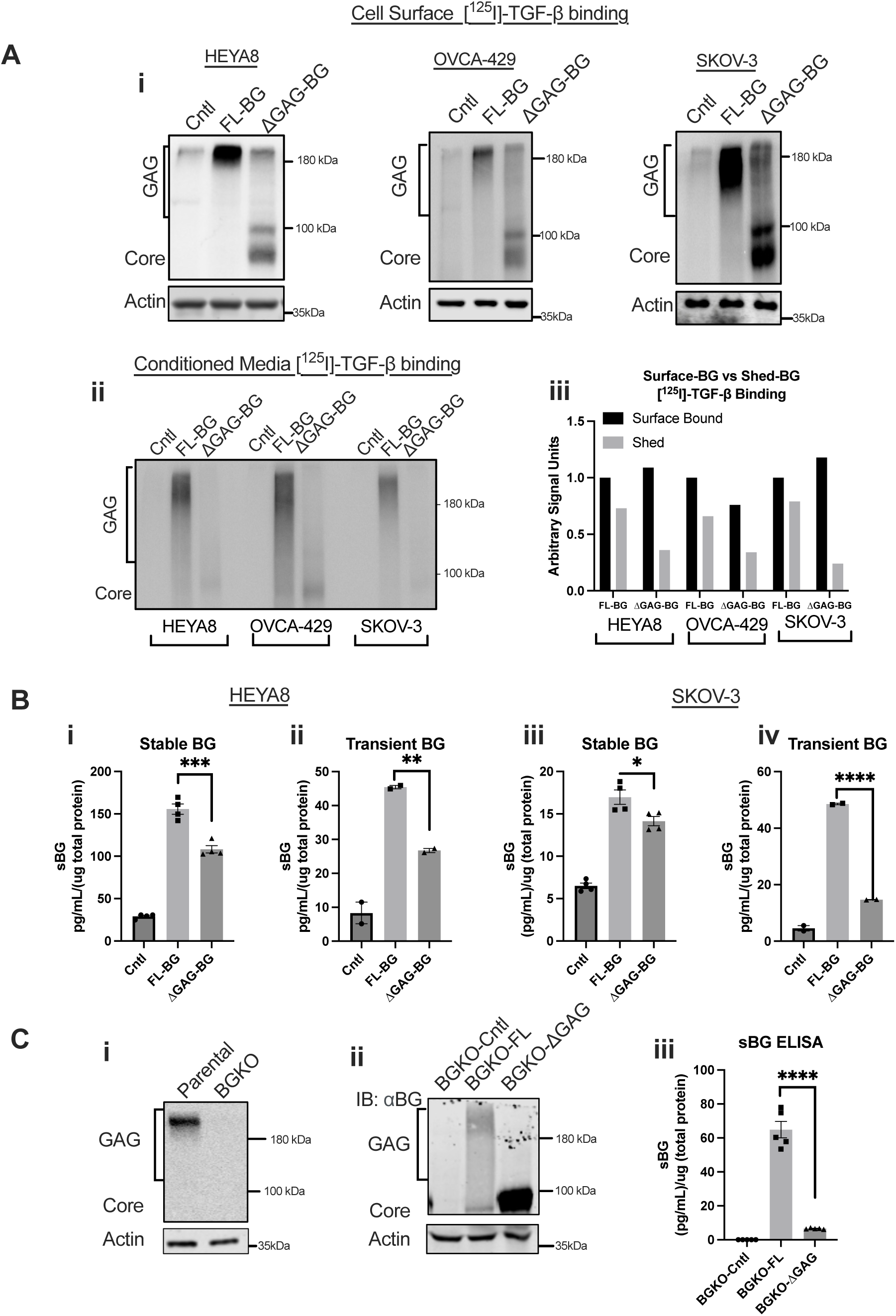
Glycosaminoglycan-modified betaglycan (BG) sheds more than unmodified BG. A. i) Autoradiograph of samples after [^125^ I]-TGF-β1 binding and crosslinking of either (i) total cell lysates or (ii) from conditioned media of indicated cells with stable (HEYA8, SKOV-3) or transient (OVCA-429) expression of BG (control-vector, FL-BG, ΔGAG-BG) and (iii) quantification of the cell surface binding autoradiograph to the conditioned media binding autoradiograph of indicated cells expressing FL-BG or ΔGAG-BG. B) BG-ELISA from the conditioned media of indicated HEYA8 (i, ii) and SKOV-3 (iii, iv) cells. The concentration of shed-BG normalized to the total protein concentration of the cells is plotted by Mean ± SEM, (n = 4 for stable and 2 for transient). *p < 0.05; **p <0.01; ***p < 0.001; ****p <0.0001, One-way ANOVA followed by unpaired t-test between FL-BG and ΔGAG-BG (HEYA8 stable, p=0.0007, transient, p=0.0018, FL-BG vs ΔGAG-BG) (SKOV-3 stable p=0.034, transient p<0.0001, FL-BG vs ΔGAG-BG). C. i) Autoradiograph of SKOV-3 parental and BG CRISPR Knockout (BGKO) cells radiolabeled with [^125^ I]- TGF-β1 followed by immunoprecipitation using anti-BG antibody. ii) Western blot of SKOV-3 BGKO cells transiently expressing FL-BG (BGKO-FL) or ΔGAG-BG (BGKO-ΔGAG) immunoblotted with anti-BG. iii) BG ELISA of conditioned media collected from the SKOV-3 BGKO cells expressing FL-BG (BGKO-FL) and ΔGAG-BG (BGKO-ΔGAG). The concentration of shed-BG normalized to the total protein concentration of the cells is plotted (Mean ± SEM, n = 5). ****p < 0.0001, One-way ANOVA followed by unpaired t-test between BGKO-FL and BGKO-ΔGAG.

To test the amount of shed BG as a result of cells expressing either FL-BG or ΔGAG-BG; conditioned media was subjected to [^125^ I]-TGF-β1 binding and crosslinking followed by immunoprecipitation using an anti-BG antibody to assess shed-BG. Shed-BG with glycosylated forms ranging from 90kDa to 250kDa were present in the conditioned media of FL-BG-expressing cells as previously reported ^21,38,58^ and seen here (**Fig 1A.ii, FL-BG labeled lanes**). However, media from ΔGAG-BG expressing cells had reduced [^125^ I]-TGF-β1 intensity compared to the cell surface [^125^ I]-TGF-β1 bound to ΔGAG-BG (**Fig 1A.ii ΔGAG-BG labeled lanes compared to Fig 1A.i (cell-surface BG [**^125^ **I]-TGF-β1 binding**). Quantification of [^125^ I]-TGF-β1 surface-bound BG and shed-BG signal units in conditioned media samples indicate that ΔGAG-BG expressing cells had less than half the levels of shed-BG compared to FL-BG expressing cells in all cell lines. (**Fig 1A.iii**).

To quantitively assess this difference in BG ectodomain shedding of FL-BG and ΔGAG- BG expressing cells, and to rule out differences in TGF-β binding in the media we used a BG ELISA to quantify differences in the amount of shed/soluble BG in two cell lines with low BG expression and either transiently or stably expressing FL and ΔGAG mutants (HEYA8 **Fig 1B.i, ii** and SKOV-3 **Fig 1B.iii, iv**). We find that FL-BG expressing HEYA8 and SKOV-3 cells showed significantly higher shed-BG as compared to ΔGAG-BG expressing HEYA8 and SKOV-3 cells regardless of whether BG was expressed stably or transiently (**Fig 1B**). In addition, to account for any endogenous expression of BG that may interfere with the shedding differences seen, we generated a CRISPR knockout of BG in SKOV-3 cells (**BGKO, Fig 1C.i, Suppl. Fig 1B, lane 3**). Knockout cells were restored with either FL-BG or ΔGAG-BG resulting in BGKO-FL or BGKO- ΔGAG cells (**Fig 1C.ii**). We find that BGKO-ΔGAG cells shed only 1/10^th^ as much as the BGKO- FL cells (65 pg/mL/(ug total protein) compared to 6 pg/mL/(ug total protein in ΔGAG-BG expressing cells (**Fig 1C.iii**). Together, these data in overexpression and knockout backgrounds demonstrate unequivocally, quantitative differences in the shedding of fully modified betaglycan as compared to unmodified BG.

### Heparan sulfate modifications preferentially enhance BG shedding

BG can present both HS and CS modifications ^20–25^ in ovarian cancer cell lines as confirmed by enzymatic digestion using Chondroitinase ABC and Heparinase III (**Suppl. Fig 2**). To assess whether HS and/or CS differentially impact shedding, we first utilized CHO-K1 as control (wild type), and CHO-677 cells that are devoid of N-acetylglucosaminyltransferase and glucuronosyltransferase activities leading to the absence of HS GAG modifications ^59^, and CHO- 745 mutant cells that are xylosyltransferase deficient leading to the absence of both HS and CS modifications ^60^. BG-FL expressed in CHO-K1 WT, 677, and 745 cells respectively presents modifications as anticipated (**Fig 2A.i**) and as described previously ^49^. Analysis of conditioned media from the CHO cells for soluble BG by ELISA revealed that WT CHO cells expressing fully GAG-modified BG shed twice as much as the single CS chain expressing 677 cells (WT = 3.3 ng/mL/(ug total protein) vs. 677 = 1.6 ng/mL/(ug total protein) (**Fig 2A.ii**). The greatest reduction was seen upon eliminating HS chains (CHO 745 cells) as compared to WT BG expressing cells (3.3ng/mL/(ug total protein) to 0.9ng/mL/(ug total protein) (**Fig 2A.ii**). While a further small reduction in shedding was seen in CHO 745 cells that lacked all modifications (**Fig 2A.i, ii third lane**), the differences between BG in CHO745 and CHO 677 were statistically significant, suggesting that eliminating HS modifications was likely the largest contributor to the shedding reduction of BG.

**Fig 2.**
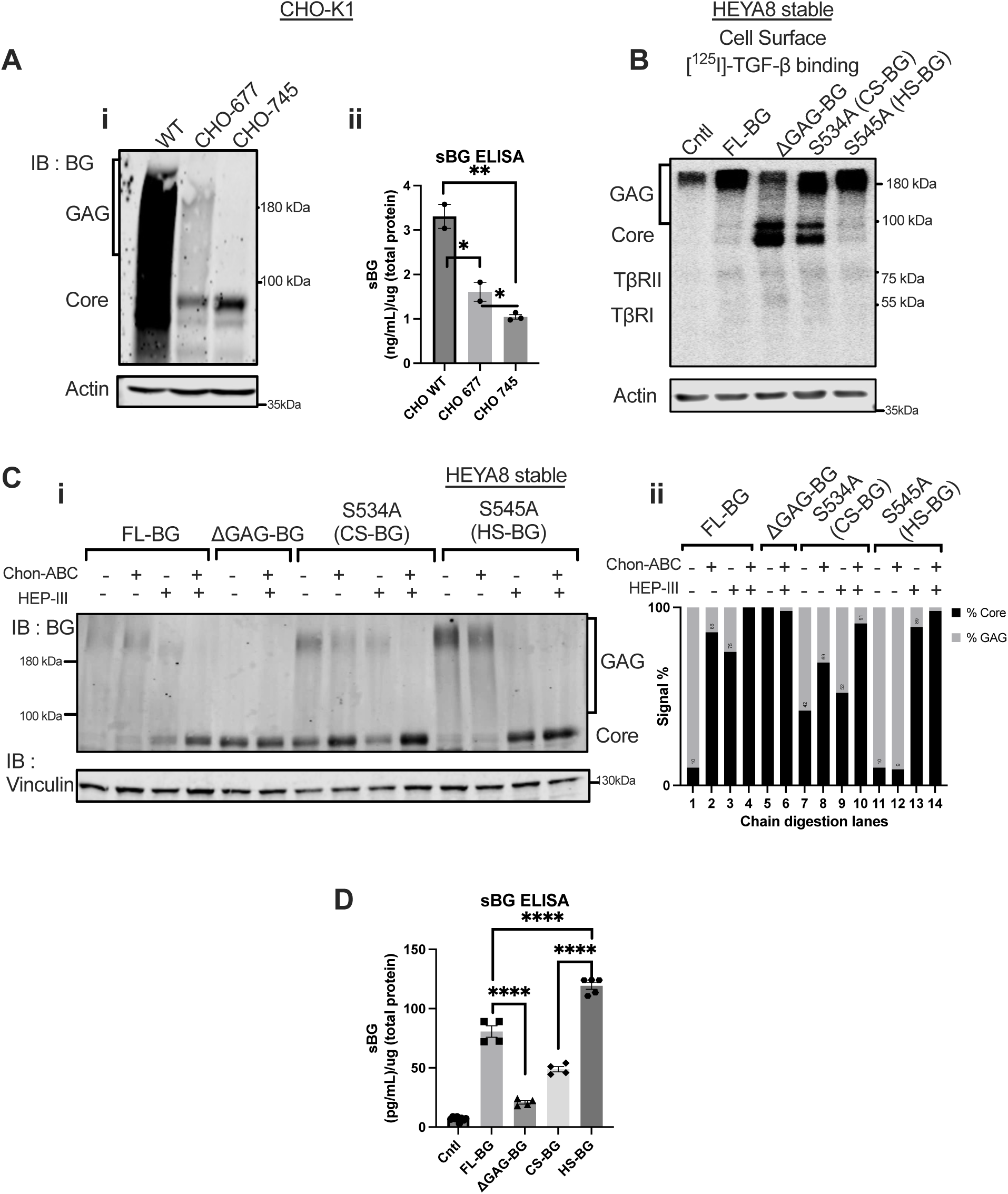
Heparan sulfate modifications of BG promote ectodomain shedding. A. i) Western blot of BG in CHO-K1 WT, 677, and 745 cells expressing FL-BG. ii) ELISA of BG from conditioned media collected from CHO-K1 WT,677, and 745 cells expressing FL-BG. The concentration of shed-BG normalized to the total protein concentration of the cells is plotted by Mean ± SEM, (n = 2) *p < 0.05; One-way ANOVA followed by unpaired t-tests between CHO WT vs CHO 677, CHO WT vs CHO 745, and CHO 677 vs CHO 745 (p=0.039 WT vs 677, p=0.0018 WT vs. 745, p=0.0456, 677 vs. 745). B. i) Autoradiograph of samples after [^125^ I]-TGF-β1 binding and crosslinking of cell surface receptors followed by immunoprecipitation using anti-BG antibody in indicated cells. C. i) Western blot of BG from indicated HEYA8 cells after enzymatic treatment using chondroitinase-ABC (0.4U) and/or heparinase-III (50ng/mL). ii) Quantification of western blot from i. The signal intensity of the BG-core band at 90kDa was compared to the intensity of the larger GAG-modified BG intensity spanning 100kDA to 250kDa. Graph is plotted as %GAG to % core signal intensity normalized to the actin levels. D. BG ELISA of the conditioned media collected from indicated HEYA8 cells. The concentration of shed-BG normalized to the total protein concentration of the cells is plotted by Mean ± SEM, (n = independent trials for Cntl = 8, FL-BG = 4, ΔGAG- BG = 4, S534A (CS-BG) = 4, S545A (HS-BG) = 5).; ****p <0.0001, One-way ANOVA followed by unpaired t-test. p = <0.0001, FL-BG vs ΔGAG-BG, p < 0.0001, CS-BG vs. HS-BG, p=0.0002, FL-BG vs. HS-BG).

To specifically test the effects of such HS modifications on BG further, we generated HEYA8 stable cell lines expressing BG with either predominantly HS or predominantly CS forms of GAG chains (**Fig 2B, 2C**). This was accomplished by generating single point mutations of S545A or S534A followed by expression of the mutants in HEYA8 cells. S534A was previously reported to be modified by both CS and HS with a preference for CS modifications ^44,46,61^ and S545 was previously reported to be modified primarily by HS GAG chains ^2,44,49,52^. [^125^ I]-TGF-β1 cell surface binding and crosslinking confirmed the cell surface availability and TGF-β binding capability of the BG mutants (**Fig 2B**). Chondroitinase ABC and Heparinase III digestion of BG GAG chains confirmed the presence of both CS and HS modifications on S534A (**Fig 2C.i lanes 7-10, 2Cii lanes 7-10**) with complete loss of modifications in S545A after heparinase III digestion (**Fig 2C.i, lane 13, 2C.ii lane 13**). BG ELISA of the shed BG (sBG) in the conditioned media of the BG-expressing cells revealed that cells expressing HS only (S545A) BG shed 1.5 times more than cells expressing FL BG (119pg/mL versus 80pg/mL respective) (**Fig 2D**) and 2.5 times higher than cells expressing S534A (CS modifications alone 49pg/mL versus S545A-HS BG at 119pg/mL) (**Fig 2D**). The least amount of shedding was seen in the ΔGAG cells as seen in (**Fig 1 and Fig 2D**). The use of the individual mutants along with the CHO cell lines together indicates that HS modifications on BG facilitate shedding to a greater extent than CS modifications with a significant reduction seen upon loss of both modifications.

### GAG modifications on BG are critical for fine-tuning TGF-β cellular signaling and cell migration responses

Soluble/shed betaglycan has been shown to sequester TGF-βs’ ^19,44,48^. Hence, we tested if ΔGAG-BG influences signaling and phenotypic TGF-β responses. Prior studies indicate maximum phosphorylation of SMAD2/3 by 30 minutes ^62,63^, with OVCA cells showing similar kinetics of phosphorylation of SMAD2/3 at 30 minutes (**Suppl. Fig 3A**). Hence using this duration (30 minutes), we tested a dose range of TGF-β concentrations required to phosphorylate SMAD2/3. Expression of BG-FL in BGKO cells led to a 40 – 55% reduction in TGF-β1-induced phosphorylation of SMAD2/3, compared to control (BGKO-Cntl) cells (**Figure 3A.i SKOV-3**). Similarly, compared to control HEYA8 cells, BG-FL expressing cells suppressed phosphorylation of SMAD2/3 in response to TGF-β1 treatment by 40% at the lowest dose of 25pM and continued to suppress by 60% in higher doses of 100pM as well (**Fig 3B.i**). In response to TGF-β2, FL- BG suppressed SMAD2/3 phosphorylation significantly at both low and high doses depending on the cell line (**Fig 3A.ii, Fig 3B.ii, Suppl. Fig 3D.ii**). Expression of ΔGAG-BG however showed a complete lack of the suppression of TGF-β1 signaling seen in FL-BG cells consistently across all cell lines (**Fig 3A.i, B.i, Suppl. Fig 3D.i**). In the case of TGF-β2 as well, ΔGAG-BG did not suppress TGF-β2 induced SMAD2/3 signaling (**Fig 3A.ii, 3B.ii, Suppl. Fig3 D.ii**). Notably, ΔGAG-BG expressing cells in BGKO cells (BGKO-ΔGAG), led to a further increase in TGF-β2 signaling even as compared to control cells (**Fig3A.ii**). These data together suggest that loss of GAG modifications on BG led to a complete failure to suppress TGF-β1 signaling and an enhancement of TGF-β2 signaling.

**Fig 3.**
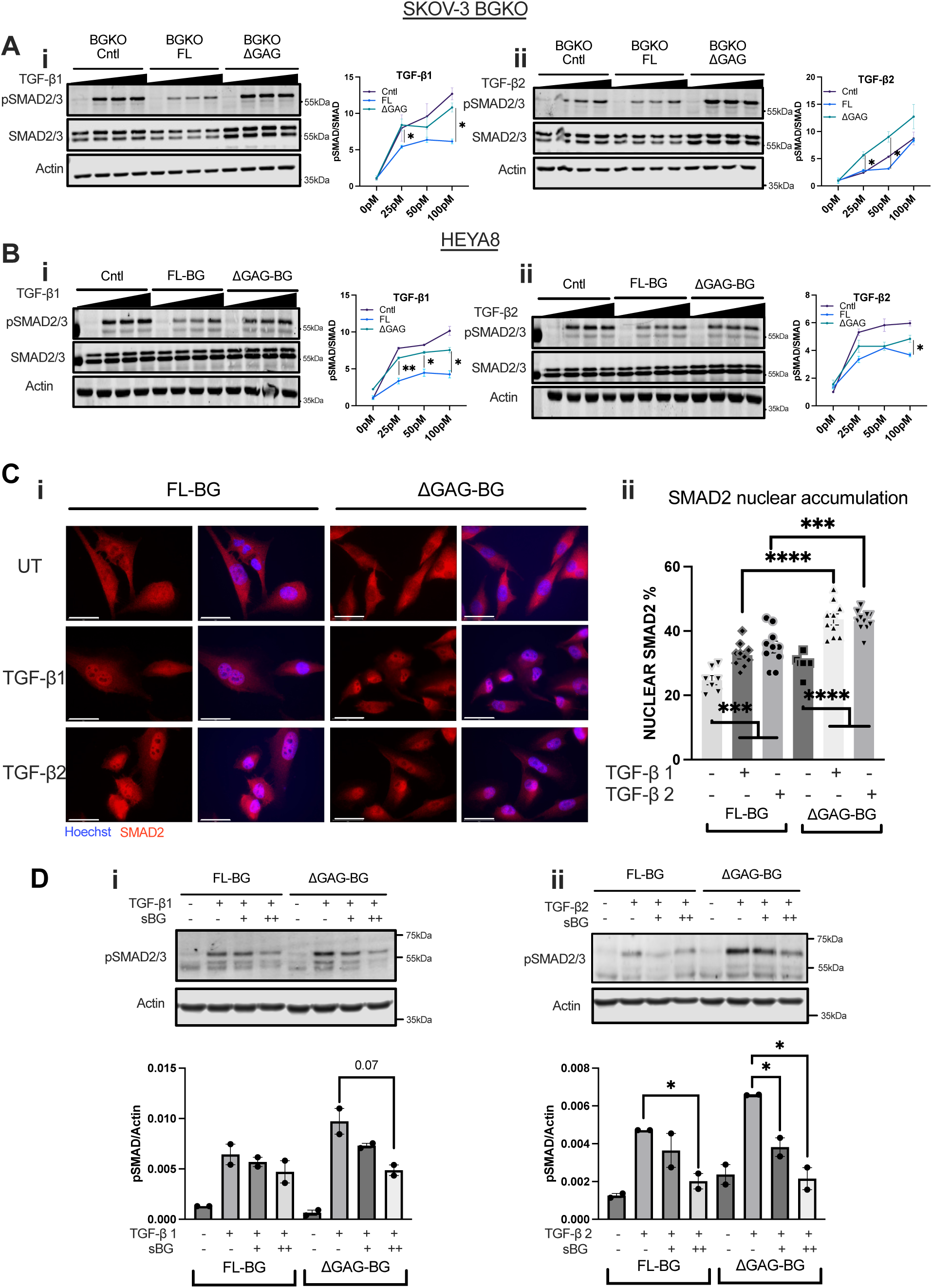
Suppression of TGF-β signaling by BG is dependent on the presence of its GAG chains. A. Western blot and signal quantification of SMAD2/3 phosphorylation in SKOV-3 BGKO cells expressing either control vector (BGKO-Cntl), FL-BG (BGKO-FL) or ΔGAG-BG (BGKO-ΔGAG) treated with i) increasing doses of TGF-β1 or ii) TGF-β2 (25pM to 100pM). All signaling quantifications were performed by normalization of phospho-SMAD2/3 to the total SMAD2/3 signal. Phospho-SMAD signal normalized to SMAD signal is plotted by Mean ± SEM, (n = 3 combined trials). *p < 0.05, unpaired t-test between FL-BG and ΔGAG-BG within the same TGF-β treatment dose. TGF-β1 at 25pM, p=0.0238, 100pM, p=0.0214, FL- BG vs. ΔGAG-BG. TGF-β2 at 25M, p=0.0167, 50pM TGF-β2, p = 0.0185, FL-BG vs. ΔGAG-BG. B. Western blot and signal quantification of SMAD2/3 phosphorylation in HEYA8 cells expressing either control, FL- BG, or ΔGAG-BG, treated with increasing doses of i) TGF-β1 or ii) TGF-β2 (25pM to 100pM). All signaling quantifications were performed by normalization of phospho-SMAD2/3 to the total SMAD2/3 signal. Phospho-SMAD signal normalized to SMAD signal is plotted by Mean ± SEM, (n = 3 combined trials). *p < 0.05, **p <0.01, unpaired t-test between FL-BG and ΔGAG-BG within the same TGF-β treatment dose. TGF-β1 at 25pM, p=0.0093, 50pM, p=0.0.193, 100pM p=0.0174, FL-BG vs. ΔGAG-BG. TGF-β2 at 100pM, p=0.0445, FL-BG vs. ΔGAG-BG . C. i) Representative immunofluorescence images of SMAD2 or nuclei (Hoechst) in response to 25pM of TGF-β1 or TGF-β2, in HEYA8 cells. Scale Bar = 50μm. ii) Quantification of nuclear accumulation of SMAD2 using cell profiler. The ratio of nuclear SMAD2 compared to total cellular SMAD2 is presented. Mean ± SEM, (n=7 replicates). ****p<0.0001, One-way ANOVA followed by unpaired t-test. (p <0.0001, TGF-β1, and p = 0.0007, TGF-β2, FL-BG vs ΔGAG-BG.) D. Western blot of phospho- SMAD2/3 in FL-BG and ΔGAG-BG expressing HEYA8 cells treated with 25pM i) TGF-β1 or ii) TGF-β2 either alone or in combination with (+) 200pg/mL to (++) 400pg/mL of recombinant sol-BG. Phospho-SMAD signal normalized to actin signal is plotted by Mean ± SEM, (n = 2). *p < 0.05; **p<0.05, unpaired t-test between TGF-β treated compared to TGF-β + sBG combination. (p = 0.0217, FL-BG TGF-β2 treated vs. TGF-β2 + 400pg/mL sol-BG, and p = 0.0302, ΔGAG-BG TGF-β2 treated vs. TGF-β2 + 200pg/mLsol-BG, p = 0.0171, ΔGAG-BG TGF-β2 treated vs. TGF-β2 + 400pg/mLsol-BG)

To examine if increased SMAD2/3 phosphorylation seen in ΔGAG-BG expressing cells led to SMAD2/3 nuclear accumulation, we examined SMAD2 localization in stable HEYA8 cell lines expressing either FL-BG or ΔGAG-BG. SMAD2 alone was chosen for immunostaining compared to both SMAD2/3 as SMAD3 can accumulate in the nucleus regardless of the level of phosphorylation of receptors by TGF-βs. We used 25pM of TGF-βs 1 and 2, the minimum dose required to detect differences in SMAD2/3 phosphorylation in (**Fig. 3A, 3B**), and allowed for 1hr of nuclear accumulation of SMAD2. A 1-hour time point was chosen as prior studies indicate a minimum of 45 minutes for maximal SMAD2 retention in the nucleus ^64,65^. In the absence of exogenous ligands, FL-BG and ΔGAG-BG expressing cells showed 25% and 30% of total SMAD2 respectively in the population to be in the nucleus (**Fig 3C.ii**). In FL-BG-expressing cells, TGF- β1 or TGF-β2 treatment increased nuclear SMAD2 marginally, to 32% and 35% respectively. This contrasts with control vector-expressing cells that showed 35% nuclear SMAD2 accumulation at a steady state (**Suppl. Fig 3C**) with both TGF-β1 and 2 treatments increasing the nuclear SMAD2 accumulation to 45% and 41% (**Suppl. Fig 3C**). However, in ΔGAG-BG cells, both TGF-β1 or TGF-β2 treatment increased nuclear SMAD2, to to 43% which was significantly higher than FL-BG cells treated with TGF-β1 or TGF-β2 (**Fig 3C.ii**) and to a similar extent as control vector cells (**Suppl. Fig 3C**). This result reinforces the TGF-β1,2 signaling suppression role of FL-BG that was abrogated by ΔGAG-BG cells.

We next determined if shedding of BG within 30 minutes of TGF-β treatment was responsible for the reduction of TGF-β signaling in FL-BG expressing cells. We first assessed the amount of shed betaglycan in the conditioned media of HEYA8 Control Vector, FL-BG, and ΔGAG-BG stably expressing cells in a shorter, time frame of 1 minute to 30 minutes. We find that by 30 minutes at steady state, FL-BG media contained 170 pg/mL, whereas ΔGAG-BG expressing cells shed 50 pg/mL (**Suppl. Fig 3E**). Notably, 200pg/mL of exogenous recombinant sol-BG (similar concentration of shed-BG at 30 minutes in FL-BG media) was sufficient to reduce TGF-β1,2 signaling in the ΔGAG-BG expressing cells (**Fig 3D.i.ii**). These data indicate that increased TGF-β1 and TGF-β2 signaling in ΔGAG-BG cells could be fully reduced to FL-BG levels by increasing the levels of soluble BG.

BG has been in several prior studies shown to be a strong regulator of both TGF-β dependent and independent cell migration and invasion responses ^13,54,66^. Since ΔGAG-BG sheds less and does not suppress TGF-β signaling, we tested the impact of FL-BG and ΔGAG-BG expression on cellular motility and invasion of ovarian cancer cells and the effect of TGF-β signaling on the same. We find that FL-BG lowered invasion in HEYA8 cells (**Fig 4A.ii**) that reached significance in CRISPR SKOV-3 BGKO cells which had FL-BG expression restored (BGKO-FL, **Fig 4A.iv**). In contrast, ΔGAG-BG expression in both cell lines (HEYA8 and SKOV- 3 BGKO cells) showed an increase in cellular invasion as compared to FL-BG mutants (2x in HEYA8 stable and transient, and a 1.4X increase in SKOV-3 BGKO, as well as in SKOV-3 Stable BG expressing cells, **Fig 4A.ii,4A.iv, Suppl. 4A.ii, 4B.ii**). Next, we sought to test whether exogenous shed-BG impacts cellular motility and invasion in FL-BG and ΔGAG-BG-expressing cells. We find that exogenous sBG suppressed invasion of ΔGAG expressing cells by 60%, compared to untreated ΔGAG cells (**Fig 4B**), whereas FL-BG expressing cells were not affected by exogenous shed-BG, indicating that the increased invasion in ΔGAG-BG expressing cells was, in part, due to reduced shedding of BG in ΔGAG-BG expressing cells. Moreover, to identify, whether the increased invasiveness of ΔGAG-BG expressing cells compared to FL-BG expressing cells is due to increased TGF-β signaling, we inhibited TGF-β signaling using A83-01, a small molecular inhibitor of ALK4,5,7^67^ (**Suppl. Fig 4C**) and tested for cellular invasion. A83-01 treatment led to a 50% reduction in invasion of ΔGAG-BG cells compared to untreated cells (**Fig 4C.ii**). No significant effect of the inhibitor was seen in either the control vector or FL-BG- expressing cells (**Fig4C.ii**). These data indicate that the reduced shedding and increased TGF-β signaling in the BG-ΔGAG cells is a direct contributor to increased invasion in cells that express unmodified ΔGAG-BG.

**Fig 4.**
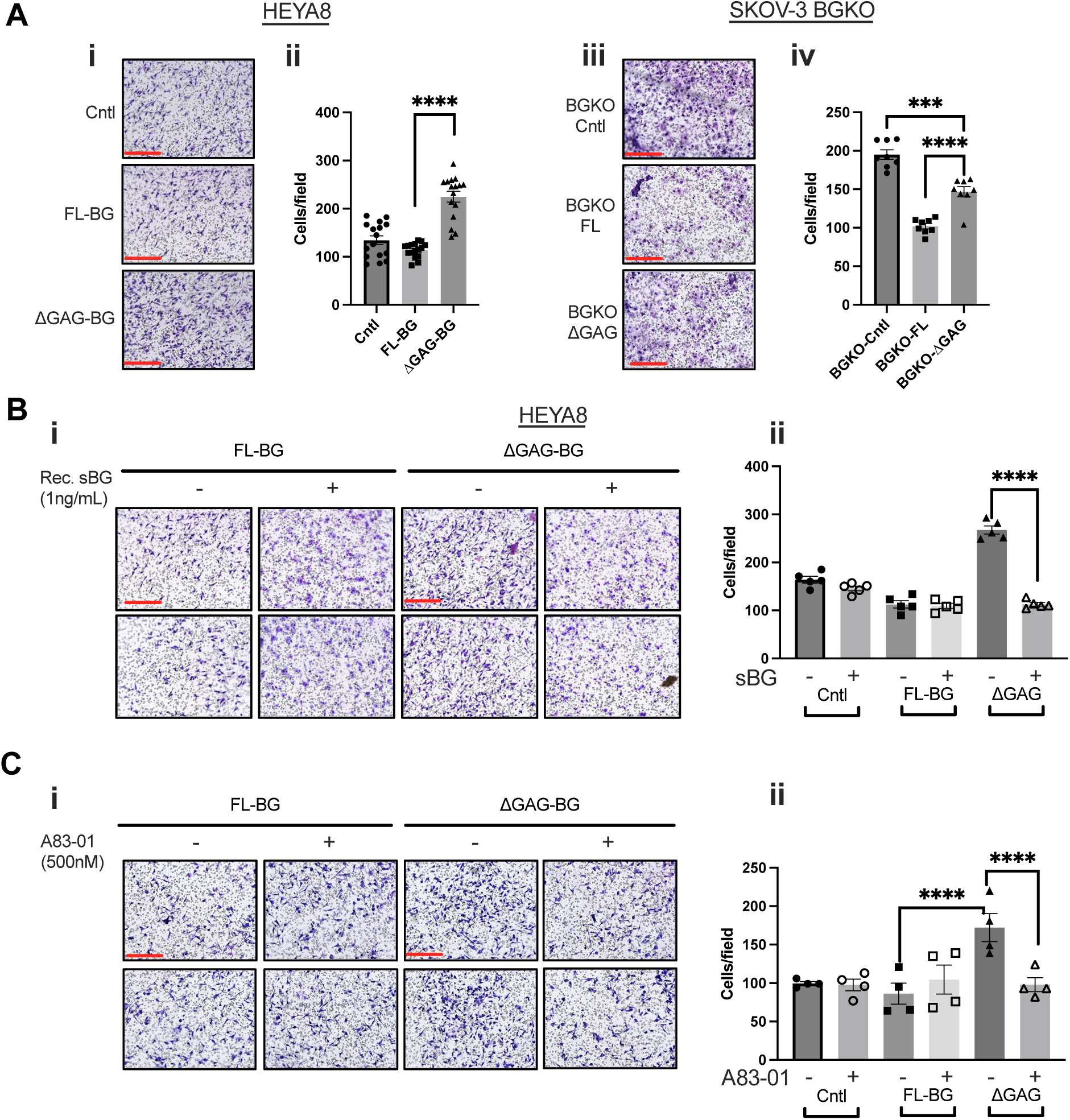
Unmodified BG (ΔGAG) promotes invasion in a TGF-β signaling-dependent manner. A. i) Representative images of invasion through Matrigel-coated Boyden-transwell inserts of HEYA8 cells expressing control vector, FL-BG or ΔGAG-BG scale bar = 275mm. ii) Quantification from (i). iii) Representative images of invasion through Matrigel-coated Boyden-transwell inserts of SKOV-3 BGKO cells as indicated. iv) Quantification of BGKO from (iii). Mean ± SEM are plotted, (HEYA8 Stable n = 16, SKOV-3 BGKO, BGKO-FL, and BGKO-ΔGAG, n = 8) ***p < 0.001; ****p <0.0001, One-way ANOVA followed by unpaired t-test between FL-BG and ΔGAG-BG. (p <0.0001, HEYA8 Stable FL-BG vs ΔGAG- BG, p <0.0001, HEYA8 Transient FL-BG vs ΔGAG-BG, p<0.0001, SKOV-3 BGKO-FL vs. BGKO-ΔGAG). B. i) Representative images of invasion through Matrigel-coated Boyden-transwell inserts of HEYA8 FL-BG and ΔGAG-BG cells treated with soluble recombinant BG at 1ng/mL in the top chamber. Scale Bar = 275μm. ii) Quantification of the invaded cells from (i). Mean ± SEM are plotted, (n = 5), ****p <0.0001, One-way ANOVA followed by unpaired t-test between sBG untreated to sBG treated groups. (p<0.0001 ΔGAG-BG untreated vs. ΔGAG-BG, sBG treated). C. i) Representative images of invasion through Matrigel-coated Boyden-transwell inserts of HEYA8 FL-BG and ΔGAG-BG cells treated with 500nM of A83-01 in the top chamber. ii) Quantification of invaded cells from (i). Mean ± SEM were plotted, (n = 4), ****p <0.0001, One-way ANOVA followed by unpaired t-test between untreated compared to A83-01 treated groups. (p<0.0001 ΔGAG-BG untreated vs. ΔGAG-BG, A83-01 treated)

### GAG-dependent BG ectodomain shedding is negatively regulated by TIMP3

We sought to delineate a mechanism for enhanced HS modification-dependent BG ectodomain shedding. Since shedding differences were seen under steady-state growth conditions, genome- wide gene expression profiles of HEYA8 control-vector cells as compared to ΔGAG, S534A-CS BG, and S545A-HS BG were compared using transcriptomics from cells under steady-state and regular growth media conditions. Single-chain mutants were tested instead of FL-BG mutants to differentiate chain-specific responses rather than a heterogeneous mixture of CS, HS, and ΔGAG that are present in FL-BG-expressing cells. We focused on differentially expressed genes that belonged to either MMP or ECM-related genes to identify potential regulators of shedding. Our analysis revealed only a small subset of genes (n = 9: *ADAMTS17*, *TMPRSS11CP*, *MMP3*, *TIMP3*, *PRSS48*, *PRSS36*, *ADAM8*, *ADAMTS3*, *MMP23B*) that were differentially expressed when comparing HS-BG and CS-BG against ΔGAG-BG expressing cells that belonged to either MMP or ECM-related genes (**Suppl. Table1**). Most notably, within the small subset of altered genes at steady state, tissue inhibitor of metalloproteinase-3 (*TIMP3*) was found to be most expressed in ΔGAG-BG as compared to both S534A CS or S545A HS-BG mutant expressing cells (p< 0.05.) (**Fig 5A.i**). Interestingly, we also noted that BG/*TGFBR3* levels were significantly higher in modified (single chain) expressing cells as compared to ΔGAG-expressing cells in the transcriptomics (**Suppl. Table 1**), however, this increase in *TGFBR3* mRNA was not observed at the protein level as determined by [^125^ I]-TGF-β1 binding and crosslinking (**Fig 2B**). We focused on TIMP3 changes and used semi-quantitative qRT-PCR and western blotting to extend the findings of the transcriptomics on TIMP3 levels to additional cell lines and in cells expressing either ΔGAG-BG or FL BG. We find that *TIMP3*-RNA was significantly higher in ΔGAG-BG cells as compared to FL BG cells in both HEYA8 and SKOV-3 cells (**Suppl. Fig 5A.i, ii**). TIMP3 protein levels were also higher in SKOV-3 ΔGAG-BG (**Suppl. Fig 5B**), but were not detectable in HEYA8 cells. Next, to test a direct role for TIMP3 and its inhibitory effects on BG ectodomain shedding, we utilized siRNAs to target TIMP3 in two cell lines (HEYA8 and SKOV-3) expressing control vector, FL-BG, and ΔGAG-BG. siRNAs to TIMP3 compared to scramble vector led to lowered TIMP3 (70-80% reduction) expression (**Fig 5B.i, iii**). BG ELISA of conditioned media from scramble-vector cells or siTIMP3 cells revealed a 51% and 28% increase in shed-BG from the ΔGAG-BG expressing HEYA8 and SKOV-3 cells respectively upon knockdown of TIMP3 (**Fig 5B.ii, iv**). Our findings thus demonstrate TIMP3 as a regulator of BG ectodomain shedding, that is dependent on the GAG chain-mediated steady-state differences in TIMP3 levels.

**Fig 5.**
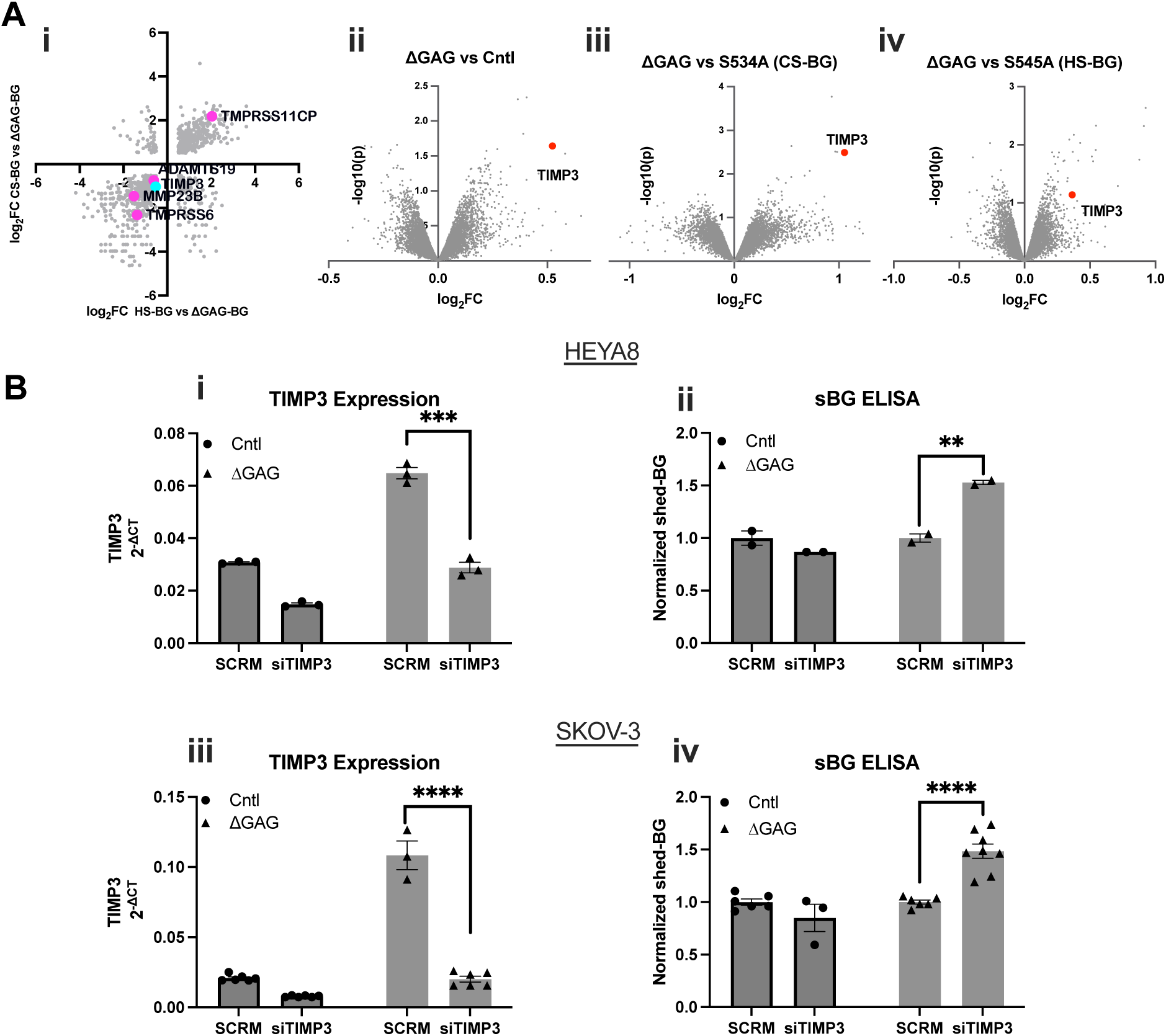
TIMP3 negatively regulates glycosaminoglycan-dependent BG ectodomain shedding. A. i) Scatter plot of the differentially expressed genes from transcriptomics of HS-BG and ΔGAG-BG plotted against CS-BG and ΔGAG-BG comparison groups. All protease-related genes are highlighted. ii) Volcano plots of the differentially expressed genes between ΔGAG-BG vs. Control-vector, (iii) S534A (CS-BG) vs. ΔGAG-BG, and (iv) S545A (HS-BG) vs. ΔGAG-BG. (*TIMP3*; ΔGAG vs Cntl, log2FC =0.524, -log(10)p = 1.643, ΔGAG vs CS-BG, log2FC =1.054, -log(10)p = 2.493, ΔGAG vs HS-BG, log2FC =0.364, -log(10)p = 1.1411.). B. i, iii) Semi-qRT-PCR of *TIMP3* and ii, iv) ELISA for sol-BG in indicated cells normalized to the scramble control cells. Mean ± SEM (n=3 qRT-PCR), *p < 0.05; **p <0.01; ***p < 0.001; ****p <0.0001, One-way ANOVA followed by unpaired t-test between scramble vector treated cells to siTIMP3 treated cells. (HEYA8 p = 0.0003 ΔGAG-BG SCRM vs siTIMP3, SKOV-3 p < 0.0001, ΔGAG-BG SCRM compared to siTIMP3), for ELISA in (Bii,iv) n=2 HEYA8, n=6 SKOV-3, p = 0.0068, ΔGAG-BG SCRM vs siTIMP3 in HEYA8 and p = <0.0001, ΔGAG scrm vs. siTIMP3 in SKOV-3.

### Shed BG is found in patient ascites fluid and correlates with patient outcomes and TGF-β responses

To assess the clinical significance of shed-BG we assessed the levels and type of BG (if any) in the ascites fluid of ovarian cancer patients. For this, acellularized ascites fluid (AF) of advanced OVCA patients from three different institutions was assessed by ELISA for soluble BG to account for any differences associated with the collection and banking of samples (Duke, Penn State, and UAB). A total of 60 samples were tested. Individual patient AF information as well as staging, survival, and histology information can be found in (**Suppl. Table 2**). We find that the range of sol-BG is between 96pg/mL to 2600pg/mL (**Suppl. Fig 6A**). The average concentration of sol- BG across all repositories was 993pg/mL, with a median of 915pg/mL (**Suppl. Fig 6A**), suggesting that potential differences in banking the fluid and age of fluid did not impact the detection of sol- BG. We thus combined the data and found that Stage 1 patients contained a mean of 854±184 pg/mL sBG, and Stage 2 had a mean of 1128±90pg/mL. The highest variability was seen in stage 3 patients, with a mean of 980±530 pg/mL and a range from 95 pg/mL at the lowest to 2632 pg/mL at the highest. Stage 4 patient samples showed the highest mean concentration of 1221 pg/mL (±463pg/mL). In comparison, serum from healthy volunteers (n=14) showed a range of 34- 100pg/mL with a median of 66.7pg/mL (**Fig 6A.i**). Although sample numbers in each stage were variable, the amount of sol-BG positively correlated with the disease stage (R^2^ = 0.91 ST1 vs ST4) (**Fig 6A.i**), indicating accumulation of sol-BG with increasing disease stage. We also find a negative correlation (r = -0.4654) between sol-BG levels and patient survival (in months) (**Fig 6B**). Stratifying the patients into low or high BG groups followed by survival analysis indicates that ‘High sol-BG’ patients had the lowest median survival (22 months) compared to ‘Low sol-BG’ patients (47.2 months) (**Fig 6C**). Since stage 3 patients account for the majority of the patient samples, we also performed a survival analysis exclusively of stage 3 patients (**Suppl. Fig 6B**). We find that median survival months for ‘High sol-BG’ patients remained at 22, and 47.2 months for ‘Low sol-BG’ patients.

**Fig 6.**
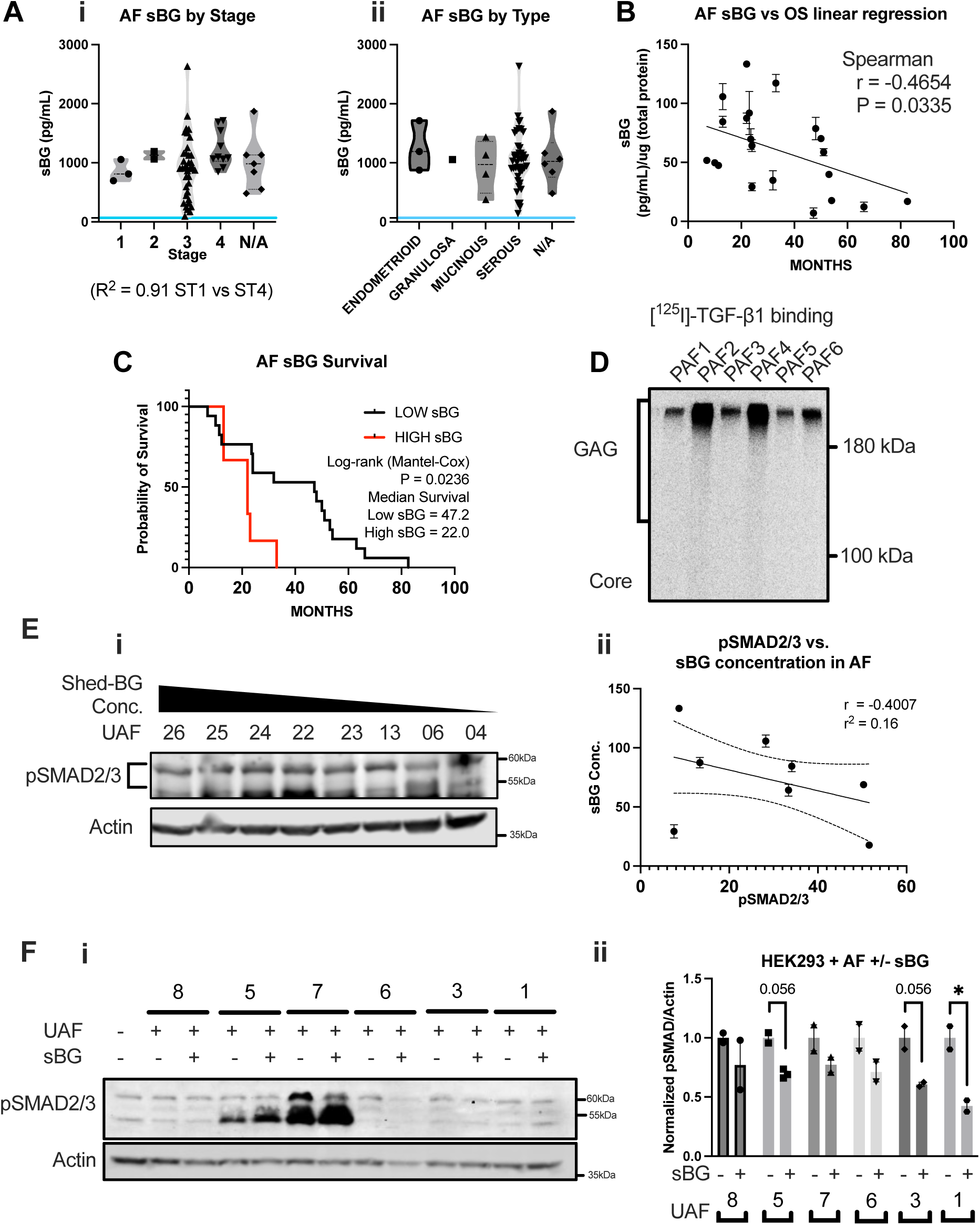
Glycosaminoglycan-modified soluble – betaglycan (sBG) in ascites fluid is associated with decreased survival and TGF-β signaling responses for ovarian cancer patients. A. i) Sol-BG concentration in AF samples plotted by i) tumor stage, and ii) tumor type. Each data point represents a single patient sample. The blue line indicates the average sol-BG concentration in the plasma of healthy volunteers. B) Correlation analysis of sol-BG concentration in the ascites fluid compared to overall survival in months. (n = 21) Spearman correlation analysis was performed, r = -0.4654, *p < 0.05; (p = 0.0335). C. Kaplan-Meier survival plot of OVCA patients stratified into low (<30th percentile) and high sol-BG (>70^th^ percentile) groups. Log-rank (Mantel-Cox) test was performed, p = 0.0236, median survival Low sol-BG = 47.2 months, High sol-BG = 22.0 months. D) Autoradiograph of patient ascites fluid following [^125^ I]-TGF-β1 binding and crosslinking and immunoprecipitation using anti-BG antibody. E. i) Western blot for SMAD2/3 phosphorylation in HEK293 cells treated with indicated OVCA patient ascites fluid. ii) Correlation graph of SMAD2/3 phosphorylation in HEK293 cells, to the sol-BG concentration in the AF used to treat HEK293 cells. Pearson correlation and simple linear regression were performed. (Pearson r = - 0.4007, R^2^ = 0.16). F. i) Western blot of SMAD2/3 phosphorylation in HEK293 cells treated with OVCA patient ascites fluid with low sol-BG concentrations (<400pg/mL), with/without 1ng/mL recombinant sol-BG. ii) Quantification of phospho-SMAD2/3 from (i) normalized to untreated samples and actin is plotted by Mean ± SEM, (n = 2), unpaired t-test between recombinant sBG untreated to treated samples. *p < 0.05; (p = 0.056 UAF5, p = 0.056 UAF3, p = 0.0343 UAF1)

We next assessed the modification status of BG in the patient ascites fluid. We find that ascites-derived BG is glycosaminoglycan modified and binds to TGF-β1 effectively (**Fig 6D**). Ascites-derived sol-BG is present with both HS and CS chains (**Suppl. Fig 6C**) similar to when expressed in vitro in cell lines (**Fig 1A, Suppl. Fig 2**), as determined by Heparinase and Chondroitinase enzymatic digestion to quantify the percent distribution of GAG chains of sol-BG in the ascites fluid (**Suppl. Fig 6C.i, ii**). To assess if sol- BG in the fluid impacted TGF-β ligand availability for cell signaling, HEK293 cells were treated with patient fluid (n= 8) for 30 minutes and analyzed for phosphorylation of SMAD2/3 (**Fig 6E.i**). AF samples were chosen based on their sBG levels ranging from the highest sBG concentration at 2600pg/mL to the lowest at 250pg/mL (**Suppl. Fig 6D, Suppl. Table 2**) We find that cells treated with UAF26, which contain the highest amount of sBG at 2600pg/ml, had the least amount of SMAD2/3 phosphorylation, compared to cells treated with UAF06, containing sol-BG at 316pg/mL, and showing highest SMAD2/3 phosphorylation. However, not all samples impacted SMAD2/3 phosphorylation based on the sol-BG levels as UAF04, with sBG concentration at 250pg/mL, did not lead to SMAD2/3 phosphorylation in HEK293 cells. A correlation analysis of the SMAD2/3 signal in HEK293 cells compared to the concentration of the sol-BG in the ascites fluid that was used to treat HEK293s showed a negative correlation between sol-BG in the patient AF and SMAD2/3 phosphorylation in HEK293 cells (r=-0.4007) (**Fig 6E.ii**). To conversely test if SMAD2/3 phosphorylation in 293 cells in response to AF could be suppressed by the addition of rec. sol- BG, we added 1ng/mL (average of sBG concentration within the patient AF samples (**Fig 6F.i**), to the AF with the lowest sol-BG levels (**Suppl. Fig 6E, Suppl. Table 2**). Combination of AF with 1ng/mL rec. sol-BG diminished the ability of the AF to stimulate phosphorylation of SMAD2/3 by 30% in 293 cells (**Fig 6F.i, ii**). These data indicate that sol-BG in advanced OVCA patient ascites fluid is glycosaminoglycan modified and negatively correlates with patient survival. Notably, AF samples with low sBG can activate TGF-β signaling in a paracrine manner more effectively than AF from patients with high sBG. These data together suggest that sol-BG could serve as a measure of TGF-β responsiveness in patients.

## Discussion

Here, we investigated the effects of glycosaminoglycan modifications on BG ectodomain shedding and TGF-β signaling and discovered that heparan sulfate modifications are crucial for betaglycan ectodomain shedding. We also found that at baseline, TIMP3 expression is higher in cells expressing BG without the GAG chains, which we find to be a regulator of betaglycan shedding, ultimately affecting TGF-β1/2 signaling responses. Importantly, we have unequivocally demonstrated for the first time a critical link between post-translational glycosaminoglycan modifications of betaglycan and TGF-β signaling responses. Our study is particularly relevant to cancer, as modified betaglycan is present in the ascites fluid of patients with ovarian cancer and can be used for predicting patient survival and TGF-β signaling responses.

The observation that GAG modifications are required for maximum BG ectodomain shedding across ovarian cancer cell lines, as well as in noncancer cells (**Fig 1&2**) suggests that this may not be a tumor cell-specific phenomenon and could occur in host tissues in the tumor environment. We also observed that the GAG-modified BG in OVCA presents predominantly with heparan sulfate chains which also appears to be a stronger driver of shedding as compared to the CS modification (**Fig 2D**). It is unclear if the CS modifications have alternate roles when on BG, such as mediating association with ECM components as has been reported for other PGs such as NG2 ^68^. Other cell surface heparan sulfate proteoglycans (HSPG), such as syndecans, perlecans, and glypicans have been shown to promote cancer pathogenesis (including ovarian) and expression changes of such HSPGs have been correlated with poorer patient outcomes ^69–71^. Additionally, aberrant regulation of heparan sulfate sulfotransferases, responsible for enzymatic modification of HS on all HS proteoglycans, has been known to affect several pathophysiological processes from inflammation, to organ development and cancer ^50,72^ suggesting that shifting the balance to a more HS-modified BG could be common in pathologies.

Sulfation patterns on proteoglycans ^50,53,73^ can influence the accessibility of proteolytic enzymes, and alter the interactions with ligands ^74^. In the case of BG, sulfation of GAG chains was shown to impact the effect of BG on WNT signaling, wherein non-sulfated GAG chains of BG stimulated Wnt signaling ^49^. BG also binds FGF2 through GAG chains ^55,57^. In neuroblastoma, neuronal differentiation occurs in a BG GAG chain-dependent manner, and complex formation with FGF receptors and BG was GAG chain-dependent ^55,57^. Highly sulfated regions of HS, also known as s-domains ^75^, have been shown to bind FGF. In addition, HS can serve as a scaffolding molecule^76^. Future studies examining the effect of GAG interacting growth factors of BG (FGF and Wnt) and the impact of sulfation on BG ectodomain shedding and signaling are warranted.

Previous reports have shown that the affinity of TGF-β1 to BG devoid of GAG chains are comparable to those of wild-type BG ^44^ leading to the long-standing view that the GAG modifications do not significantly contribute to TGF-β signaling responses. However, our findings challenge this view indirectly, as we find that GAG chains influence shedding which has an undeniable effect on the sequestration of ligands by shed-BG, thereby influencing signaling (**Fig 3D**). Indeed we anticipate that small changes in the amount of soluble betaglycan can contribute to large changes in TGF-β signaling. It is well established that the amount and duration of TGF- β signaling are both critical to responses ^77–79^. A small increase in steady-state SMAD2 nuclear accumulation was seen in untreated ΔGAG-BG-expressing cells compared to untreated FL-BG-expressing cells (**Fig 3C**). Congruently, we observed increased phopho-SMAD2/3 in ΔGAG-BG cells compared to FL-BG cells at a steady state in serum-containing media (**Suppl. Fig 3B**). Increased phosphorylation and increased nuclear accumulation of SMAD2 upon TGF-β 1&2 treatment in ΔGAG-BG cells compared to FL-BG cells support the direct influence on durable cell signaling. Our studies focused on SMAD2/3 signaling as a primary readout of TGF-βs ½. However TGF-β s can elicit signaling through other transducers, such as SMAD1/5, and crosstalk with other signaling pathways via SMAD-independent pathways such as, but not limited to, ERK, JNK, P38, PI3K, Wnt, and Hh signaling pathways ^80^ is well known. The impact of shed-BG and TGF-β superfamily growth factors, as well as crosstalk signaling pathways in OVCA and other diseases, remains to be determined.

Our studies do not rule out the alternate possibility of differences in the ability of the fully modified (FL-BG) and unmodified (ΔGAG-BG) BG to influence the stability of the cell surface receptor complexes formed between BG and TβRII, TβRI receptors for TGF-β signaling. This model was supported by a prior study ^48^, which indicated that the GAG chains of BG reduced the formation of TGF-β receptor complexes, thereby inhibiting signal transduction. Although we did not thoroughly investigate the formation of TGF-β receptor complexes between BG GAG mutants, affinity binding of [^125^ I]-TGF-β1 in BG FL and ΔGAG mutant expressing cells showed the presence of TGF-βRI and TGF-βRII when immunoprecipitated using an anti-BG antibody, suggesting that GAG-modified and unmodified BG interact with TGF-βRI and TGF-βRII. We showed a greater impact on the inhibition of TGF-β signaling by the ligand-sequestration effect of shed-BG (**Fig 3D**). In light of our findings, a more quantitative assessment of the effect of the modifications on BG on receptor homo- heterodimerization is also warranted.

We and others published several prior studies on the effect of FL-BG on reducing tumor cell motility ^5,19,26,27,54,81^ primarily via Cdc42, via interactions with β-arrestin2 ^54,82^ and the potential influence of GAG chains ^55,83^. The expression of modified BG leading to inhibition of migration and the increased migration upon expressing unmodified BG was consistent with previous reports particularly in cells with BG knockout by CRISPR (BGKO) that had restored FL-BG (BGKO-FL) expression (**Fig 4A.iv**) suggesting that even small changes in the balance between modified and unmodified BG can shift the effects on tumor cell motility.

Under normal physiological conditions, BG can be detected in plasma ^84^ and milk ^85^ and has been shown to be a neutralizing agent for TGF-β. In the context of ovarian cancer, ascitic fluid (AF) accumulation in the peritoneum enables the transcoelomic tumor spread of metastatic cells ^86^. Previous studies have shown the prognostic value of identifying components in AF to predict treatment and survival outcomes ^87,88^. AF from tumor-bearing mice and patients are known to have elevated TGF-β1^89–91^. Our observation of the elevated amounts of soluble BG being associated with worse patient outcomes in conjunction with prior studies demonstrating lower BG expression in tumors suggests that there may be additional sources of soluble BG in the ascites. We propose that sol-BG in ascites, which can dull TGF-β signaling responses (**Fig 6E**), could potentially serve as a predictor for patient response to TGF-β ligand-targeted therapies.

## Material and Methods

### Cell Lines and Reagents

#### Ovarian epithelial carcinoma cell lines

HEYA8, SKOV-3, and OVCA429 were obtained from the Duke Gynecology/Oncology Bank (Durham, NC) and ATCC (ATCC® HTB-77^™^). CHO-K1 epithelial cell lines pgsA-745 (ATCC® CRL-2242^TM^), and pgsD-677 (ATCC® CRL-2244^TM^) were obtained from ATCC (Manassas, VA). HEYA8, SK-OV-3, and OVCA429 were cultured in RPMI 1640 (ATCC® 30-2001ATCC^TM^) containing l-glutamine, 10% FBS, and 100 units of penicillin-streptomycin. CHO-K1 cell lines pgsA-745 and pgsD-677 were cultured in DMEM/F-12 medium (ATCC 30-2006) DMEM: F-12 Medium contains 2.5 mM L-glutamine, 15 mM HEPES, 0.5 mM sodium pyruvate, and 1200 mg/L sodium bicarbonate, 10% FBS, and 100 units of penicillin-streptomycin. All cell lines were maintained at 37 °C in a humidified incubator at 5% CO2, routinely checked for mycoplasma, and experiments were conducted within 3–10 passages depending on the cell line. Cell line authentication was performed at the Heflin Center for Genomic Science Core Laboratories at UAB.

#### Antibodies

Human TGF-beta RIII (catalog AF-242) was purchased from R&D Biosystems (Minneapolis, MN), Phospho-SMAD2 (Ser465/467)/SMAD3 (Ser423/425) (catalog #8828), Smad2/3 (D7G7) (catalog #8685), Smad2 (D43B4) XP^®^ Rabbit mAb (catalog #5339), TIMP3 (D74B10) Rabbit mAb (catalog #5673) were obtained from Cell Signal Technology (Danvers, MA). Actin antibody (C-2) (catalog sc-8432) was obtained from Santa Cruz Biotech (Santa Cruz, CA).

#### Other reagents

[^125^ I]-Bolton-Hunter labeled Transforming Growth Factor- β1 (Human, Recombinant) (catalog NEX267) was obtained from Perkin Elmer (Waltham, MA) This product was however discontinued in 2019. Human TGF-beta RIII DuoSet ELISA (catalog DY242), DuoSet ELISA Ancillary Kit (catalog DY008), Recombinant Human TGF-beta RIII Protein (catalog 242-R3), Heparinase III (catalog 6145-GH-010) and chondroitinase ABC (catalog No. C3667) were obtained from Sigma-Aldrich, and recombinant TGF-β1, TGF-β2, and Wnt3a were purchased from R&D Systems. A83-01 (catalog 100-1041, 72024) was purchased from Stemcell Technologies.

### Plasmid constructs, generation of stable and transient expression cell lines

#### Constructs

Betaglycan (BG) constructs were generated as described previously ^19,48,49,55,81,92,93^. FL-BG consists of HA-tagged human BG in pcDNA 3.1(+) ^48,94^. The BG-ΔGAG construct was generated by introducing double ser-ala point mutation at amino acids 534 and 545 to prevent GAG modifications ^57,95–97^. Single GAG attachment site mutants S534A (CS-BG), and S545A (HS-BG) constructs were all generated by site-directed mutagenesis (Agilent Technologies 210515). Constructs were confirmed by sequencing.

#### Stable cell lines

FL-BG, ΔGAG-BG, S534A (CS-BG), and S545A (HS-BG) constructs were cloned into a pHIV-dTomato lentiviral backbone (Plasmid #21374, Addgene, Cambridge, MA) by the Center for Targeted Therapeutics Core Facility at the University of South Carolina (Columbia, SC) followed by lentiviral particle generation. Infected cells were sorted by dTomato expression at the flow cytometry core at the University of South Carolin Flow Cytometry Core Facility or UAB Flow Cytometry and Single-Cell Core facility.

TGFBR3 CRISPR knockout cell line was generated using Origene CRISPR/CAS9 genome-wide knockout kit (GE100021). 10 MOI of CRISPR BG pCAS guide virus and 8ug/mL of polybrene were used for each infection. Puromycin selection was performed on the infected cells. Surviving cells with puromycin resistance were then isolated into a monoclonal population using a limited dilution method. Wells containing a single colony of cells were then expanded and characterized by [^125^ I]-TGF-β1 binding and crosslinking followed by immunoprecipitation using BG antibody. KO clones with no BG expression were chosen for further experiments. All KO cells were maintained with 0.5ug/mL of puromycin.

#### Transient cell lines

Adenoviral constructs for FL-BG, and ΔGAG-BG were used to transduce cells at MOI of 50 to 200 IFUs/cell, and infections were performed as previously described ^54,55,98^. Origene siRNA-27 kit was used for transient knockdown of TIMP3 expression (SR304839 Locus IF 7078). Cells were reverse-transfected using Lipofectamine-RNAiMAX with 10nM of universal negative control RNA duplex (Scrambled) compared to the cell transfected with 10nM of pooled duplexes targeting TIMP3. siRNA/Scramble vectors were transfected into HEYA8 and SKOV-3 cells using RNAiMAX. Media was refreshed and samples were collected 48-72 hours post-transfection.

### Immunoprecipitation, Western Blotting, and Immunofluorescence

Immunoprecipitation and Western blotting were performed using standard techniques ^49,82,99^. For immunoprecipitations of shed-BG and membrane-bound BG, conditioned media/cell lysed in CO-IP lysis buffer (50 mm Tris-HCl, pH 7.5, 150 mm of NaCl, 1% Nonidet P-40, 10% glycerol, 1 mm DTT, 25 mm NaF, 1 mm Na3VO4 and 1× protease inhibitor mixture (catalog No. P8340, Sigma-Aldrich)) was incubated overnight with 2.5ug of anti-human BG antibody and 30uL of Protein G-Sepharose beads in 4C with mild agitation. The next day, PGS beads were washed three times with cold PBS and re-suspended in the 2x Laemmli sample buffer. For immunofluorescence, HEYA8, Cntrl-Vector, FL-BG, and ΔGAG-BG expressing cells were seeded onto coverslips in 12-well plates at a density of 5 × 10^4^ cells/well. After 24 hours, cells were serum starved overnight and then treated with 25pM TGF-β1/2 for 1hr unless otherwise indicated. Cells were fixed in 4% paraformaldehyde and permeabilized with 0.1% TritonX-100, followed by blocking with 5% BSA in PBS for 1 hr. SMAD2 was labeled using the D43B4 CST SMAD2 antibody and incubated with an Alexa-conjugated secondary antibody (Molecular Probes, Eugene, OR). Nuclei were stained with Hoechst 33258. Coverslips were mounted using ProLong Gold Antiface mountant (Thermo Fisher catalog P36930). Immunofluorescence imaging was captured using an EVOS M7000 microscope at 60x magnification. SMAD2 localization was quantified using a Cell Profiler ^100^ pipeline.

### Crosslinking and Binding with [^125^ I]-TGF-β1

Cell surface receptors and conditioned media [^125^ I]-TGF-β1 crosslinking and binding methods have been extensively described previously ^2,44,61^. Cell surface receptor binding was conducted in a cold room to inhibit receptor internalization. For cell surface labeling, 100pM of [^125^ I]-TGF- β1 in HEPES-*KRH Buffer was used* (Final concentration. NaCl, 116 mm. KCl, 4 mm. MgCl2, 1 mm. CaCl2, 1.8 mm. Glucose, 25 mm. HEPES acid, 10 mm. Adjust pH to 7.4). For shed-BG binding, conditioned media from BG GAG mutant expressing cells were incubated in full-serum media or serum-free media, as noted in the figure legends, and were directly labeled with 200pM of [^125^ I]-TGF-β1. Washing steps were omitted in conditioned media BG binding. Cell lysate samples were lysed with 2x Laemmli sample buffer or CO-IP buffer, followed by the immunoprecipitation protocol described above. SDS-page gels were dried onto a filter paper at 80C for 2.5 hrs on a gel dryer. Dried gels on filter papers were developed onto a phosphor screen for 10 – 21 days. Imaging of the phosphor screen was performed on the GE Typhoon system. The scanned image was then analyzed using ImageQuant software.

### RNA isolation and semi-quantitative RT-PCR

Total RNA was isolated using TRIzol reagent/protocol from Invitrogen. RNA was reverse transcribed using iScript Reverse Transcription Supermix and iTaq Universal SYBR Green Supermix. Expression data were normalized to the geometric mean of housekeeping genes *RPL13A* and *HPRT1.* The qRT-PCR primers sequences used were: *RPL13A* forward: AGATGGCGGAGGTGCAG; reverse GGCCCAGCAGTACCTGTTTA, *HPRT1* forward: TGACCTTGATTTATTTTGCATACC; reverse: CGAGCAAGACGTTCAGTCCT, *TGFBR3* forward: CGTCAGGAGGCACACACTTA; reverse: CACATTTGACAGACAGGGCAAT, and *TIMP3* forward: GTGGTCAGCCTCTCTCACAC; reverse: AAGACCCTTCTTTGCCCAGG.

### ELISA

Betaglycan ELISA (DY242) from R&D systems was utilized for the majority of this study apart from the processing of Duke repository AF samples, the methodology for which is as described previously ^32,34^ and performed according to the manufacturer’s instructions to quantitatively measure BG concentration in conditioned media. Conditioned media was collected in full-serum media or serum-free media for durations noted in the figure legends. Biological duplicates of samples were collected per condition and 2-3 technical replicates per biological replicate were analyzed. The optical density of each well was measured via a Gen-5 plate reader set to 450nm with wavelength correction set to 540nm. Optical density values were used to calculate the concentration of sBG using a 4PL calculator (AAT-BIO) based on recombinant human BG standard values ran with every set of experiments.

### Trans-well invasion assay

20,000 HEYA8/SKOV-3 and 100,000 SKOV-3 BGKO cells were plated on a matrigel-coated (400 μg/mL) 8 μm trans-well filter in serum-free media. 10% FBS media was used as a chemoattractant in the bottom chamber. Apical cells were scraped off and invaded cells were fixed and stained using a Three-Step stain set from Thermo Fisher. 3-5 random images were taken per filter using a 10X objective on the EVOS M7000 microscope. Cells were counted using the ImageJ Cell-counter plugin.

### Patient Ascites

Specimens from patients diagnosed with primary ovarian cancer were collected and banked after informed consent at Duke University Medical Center, Pennsylvania State University College of Medicine (Hershey, PA), or the University of Alabama Birmingham, with approval for the study grant from the Duke University’s institutional research ethics board, Penn State College of Medicine and UAB Institutional Review Boards (IRB) respectively. All samples were previously banked frozen samples. Acellular ascites fluid was briefly spun down at 1200rpm and then the supernatant was collected. The protein concentration of each AF sample was measured using a Pierce BCA Protein Assay kit (Thermo scientific 23227) and then normalized to equal concentrations for downstream analysis.

### RNA-SEQ

RNA library preparation was performed using NEBNext Ultra II Directional RNA Library prep Kit following the manufacturer’s protocols followed by a quality check of Indexed sequences by Fastqc. Indexed sequences were trimmed using an adaptor sequence by TrimGalore-0.4.5. Read counts over annotated genes were obtained using featureCount. Differential expression analysis was performed using the bioJupies web tool ^101^. RNA-seq data have been deposited in the NCBI Gene Expression Omnibus (GEO) database (Accession number GSE237403).

### Statistics

All graphs are representative of 3-5 independent biological experiments with individual points denoting the average of each experiment unless described otherwise in figure legends. Data are expressed as Mean ± SEM. Statistical analyses were performed using GraphPad Prism 11, described in the figure legends. The difference between the two groups was assessed using a two-tailed *t*-test. Multiple group comparisons were carried out by the analysis of variance (ANOVA). Survival analysis and correlation analysis were performed using GraphPad Prism 11 using the Mantel-Cox test, and Spearman correlation test.

## Data availability

- All the sequencing data are publicly available and have been deposited at the Gene Expression Omnibus (GEO) accession number GSE237403.
- Original Western blot and microscopy images are available from the corresponding author upon request.
- This study does not report any original code
- Any additional information and data for reanalysis is available from the corresponding author upon request.

## Supporting information

- Acknowledgments
- Author contributions
- Funding and additional information
- Conflict of interest

## Acknowledgments

We would like to thank Ben Horst, Kevin Tabury, Mehri Monavarian, Resha Rajkarnikar, Eduardo Listik, Emily Page, and Liz Quintero-Macias for technical assistance and helpful discussion. We also thank the Functional Genomics Core, COBRE Center for Targeted Therapeutics at the University of South Carolina for RNA sequencing, CRISPR generation, and lentiviral preparations, and the Heflin Center for Genomic Science core laboratories at UAB for cell line authentication services. Funding for this work was provided by NIH R01CA219495 to Mythreye Karthikeyan (KM). The funders had no role in study design, data collection and analysis, decision to publish, or preparation of the manuscript.

