## Supplementary Figures for "Heparan sulfate modifications of betaglycan promote TIMP3-dependent ectodomain shedding to fine-tune TGF-β signaling"

Suppl. Fig 1

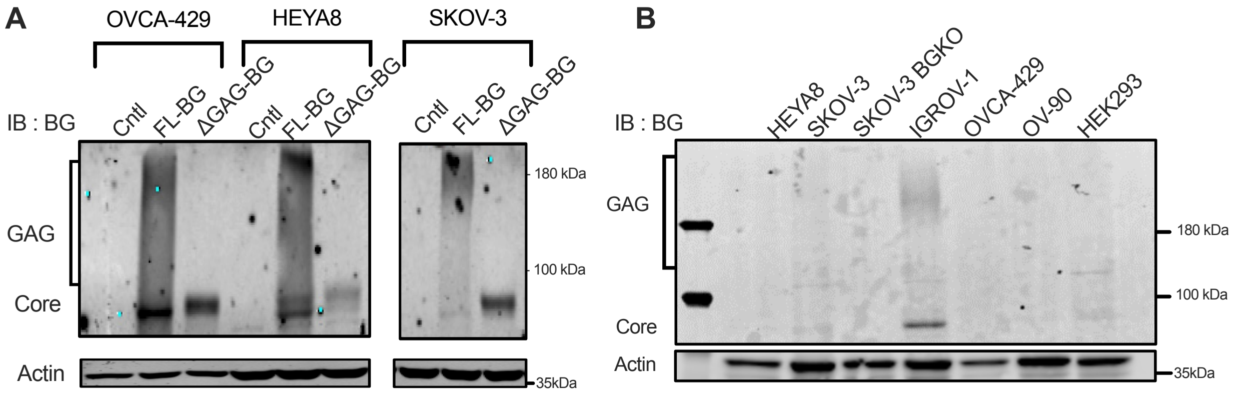


Suppl. Fig 1.

A. Western blot for BG in indicated cell lines expressing either control vector, FL-BG, or ΔGAG-BG. B. Western blot for BG in indicated cell lines at steady state.

Suppl. Fig 2

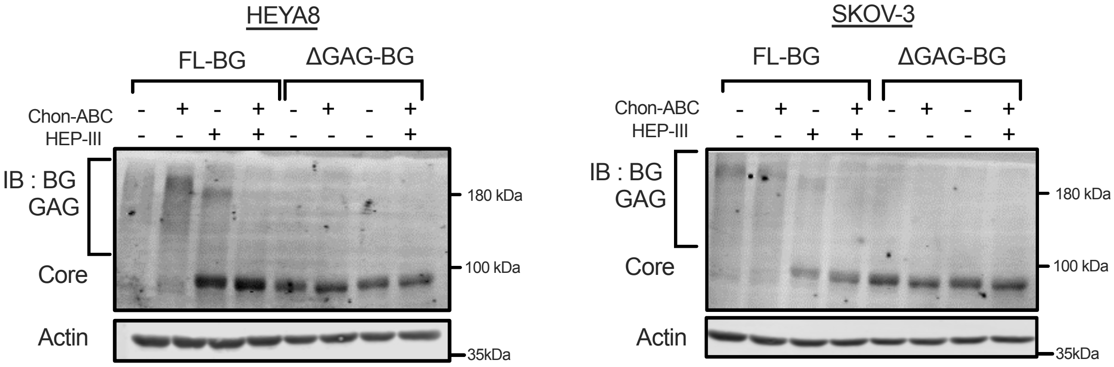


Suppl. Fig 2.

Western blot of BG in indicated cells expressing FL-BG or ΔGAG-BG subjected to enzymatic chain digestion using (0.4U) Chondroitinase and (50ng/mL) HepIII.

Suppl. Fig 3

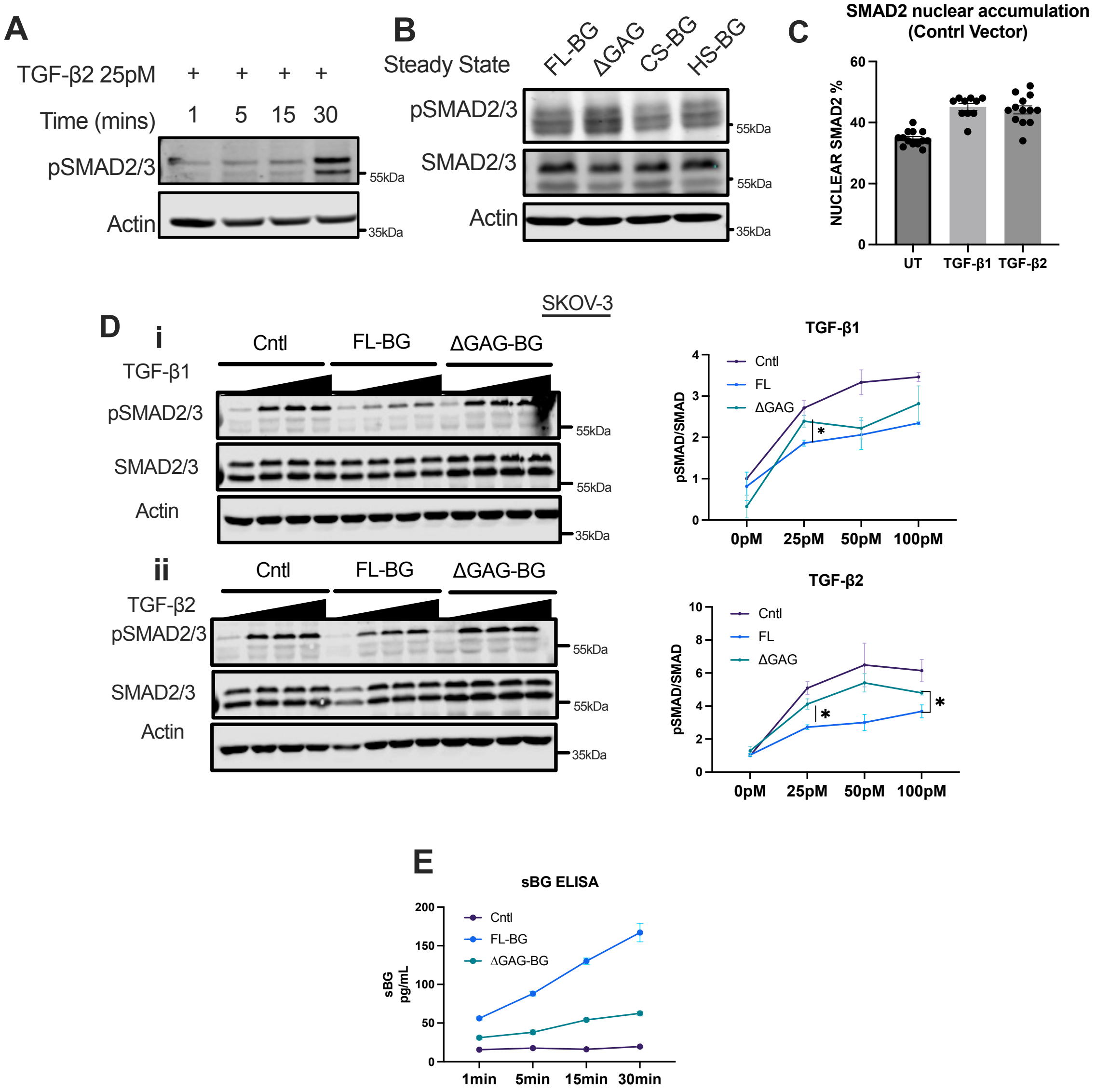


Suppl. Fig 3.

A. Western blot of phospho-SMAD2/3 in cells treated with 25pM TGF-β2 for increasing amounts of time. B. Western blot of phospho-SMAD2/3 in HEYA8 FL-BG, ΔGAG-BG, S534A (CS-BG), and S545A (HS-BG) expressing cells at steady state growth conditions. C. Quantification of nuclear accumulation of SMAD2 in HEYA8 control cells treated with 25pM TGF-β1 or TGF-β2. D. Western blot and signal quantification of SMAD2/3 phosphorylation in SKOV-3 cells expressing either FL-BG or ΔGAG-BG, treated with increasing doses of TGF-β1 or TGF-β2. All signaling quantifications were performed by normalizing phospho-SMAD2/3 to the total SMAD2/3 signal and plotted by Mean ± SEM, (n = 3). *p < 0.05; **p <0.01, unpaired t-test between FL-BG and ΔGAG-BG within the same TGF-β treatment dose. E. BG ELISA of the conditioned media collected from HEYA8 FL-BG and ΔGAG-BG cells over time in serum-free media. The concentration of the sol-BG is plotted in pg/mL (n=2)

Suppl. Fig 4

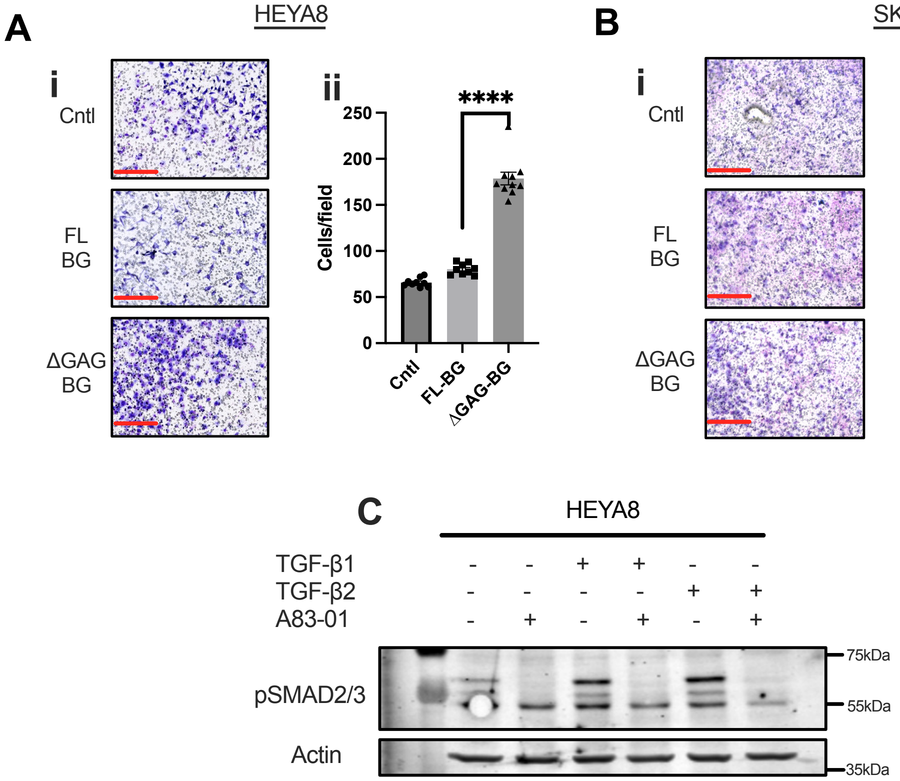


Suppl. Fig 4

A. i) Representative images of invasion through Matrigel-coated Boyden-transwell inserts of HEYA8 transient expression cells. scale bar = 275μm. ii ) Quantification of (i). Mean ± SEM was plotted, (n = 9) ****p <0.0001, One-way ANOVA followed by unpaired t-test between FL-BG and ΔGAG-BG. B. i) Representative images of invasion through Matrigel-coated Boyden-transwell inserts of SKOV-3 BG stable expression cells. Scale bar = 275μm. ii) Quantification of (i). Mean ± SEM were plotted, (n = 5) ****p <0.0001. C. Western blot of phospho-SMAD2/3 in HEYA8 cells treated with 25pM of TGF-βs 1 or 2 and in combination with 500nM of A83-01.

Suppl. Fig 5

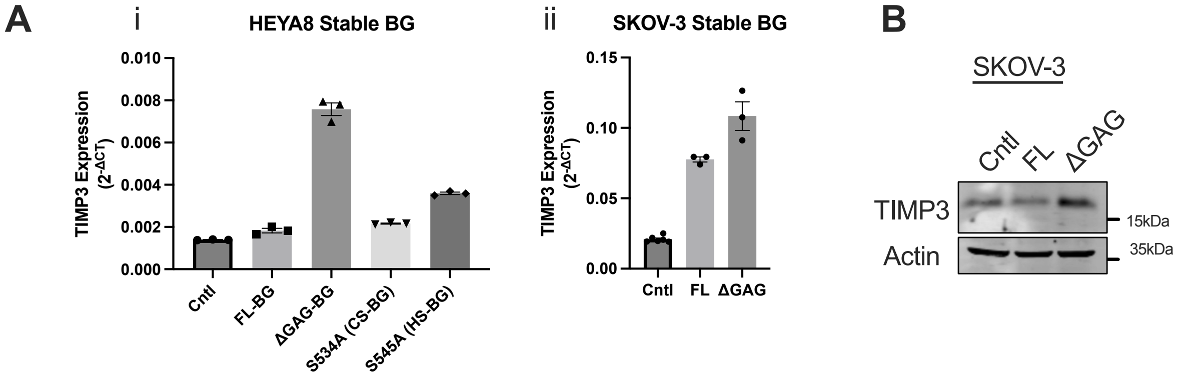


Supp. Fig 5.

A. Relative semi-qRT-PCR of *TIMP3* expression in (i) HEYA8 and (ii) SKOV-3 stable BG GAG mutants cells. (n=3). B. Western blot of TIMP3 in SKOV-3 Stable FL-BG, and ∆GAG-BG cells.

Supplementary Table 1

| S545A (HS-BG) VS ΔGAG-BG | log2FC | -log10(pval) |
|---|---|---|
| *ADAMTS17* | 0.9533 | 1.5192 |
| *TMPRSS11CP* | 2.0375 | 1.7316 |
| *MMP3* | 2.3732 | 1.9306 |
| *TIMP3* | -0.5350 | 1.9352 |
| *PRSS48* | 1.0411 | 1.4261 |
| *PRSS36* | -0.9094 | 1.6709 |
| *ADAM8* | 0.5352 | 1.4499 |
| *ADAMTS3* | -0.5870 | 1.8941 |
| *MMP23B* | -1.5254 | 1.5052 |
| *TGFBR3* | -3.1818 | 2.0212 |
| **S534A (CS-BG) vs ΔGAG-BG** | **log2FC** | **-log10(pval)** |
| *TIMP3* | -1.0342 | 5.3645 |
| *ADAMTS9* | 1.3538 | 1.6960 |
| *TMPRSS11CP* | 2.1849 | 1.6229 |
| *TMPRSS6* | -2.3199 | 2.8800 |
| *ADAM12* | 0.6949 | 1.4706 |
| *TGFBR3* | -3.4550 | 2.0981 |

Supplementary Table 1.

All protease/ECM-related genes differentially expressed in HEYA8 BG GAG mutant cells.

Suppl. Fig 6

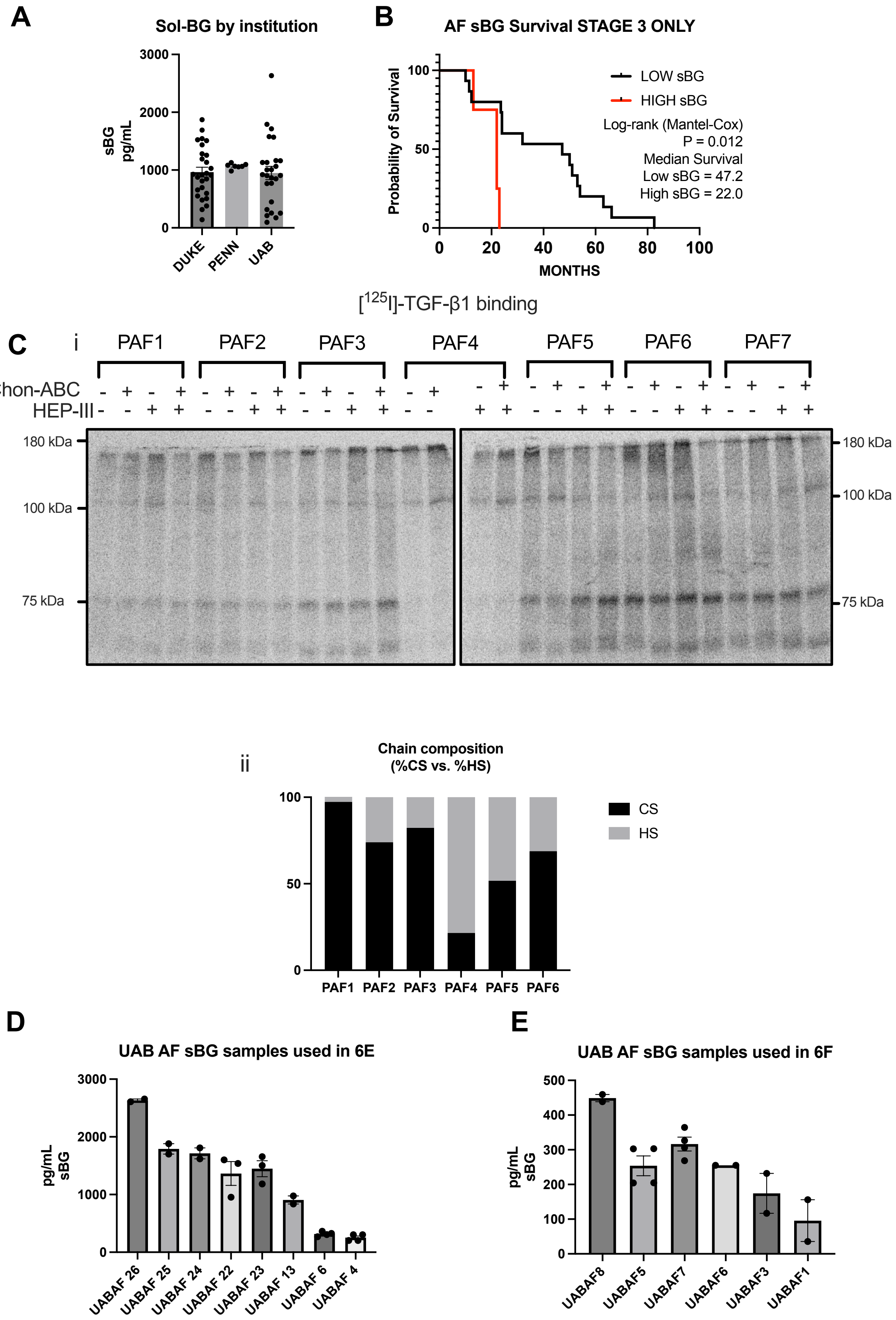
A
Sol-BG by institution
sBG pg/mL
3000
2000
1000
0
DUKE
PENN
UAB
B
AF sBG Survival STAGE 3 ONLY
Probability of Survival
100
50
0
0
20
40
60
80
100
MONTHS
LOW sBG
HIGH sBG
Log-rank (Mantel-Cox)
P = 0.012
Median Survival
Low sBG = 47.2
High sBG = 22.0
[125I]-TGF-β1 binding
C
i
PAF1
PAF2
PAF3
PAF4
PAF5
PAF6
PAF7
Chon-ABC
HEP-III
180 kDa
100 kDa
75 kDa
ii
Chain composition
(%CS vs. %HS)
100
50
0
PAF1
PAF2
PAF3
PAF4
PAF5
PAF6
CS
HS
D
UAB AF sBG samples used in 6E
pg/mL sBG
3000
2000
1000
0
UABAF 26
UABAF 25
UABAF 24
UABAF 22
UABAF 23
UABAF 13
UABAF 6
UABAF 4
E
UAB AF sBG samples used in 6F
pg/mL sBG
500
400
300
200
100
0
UABAF8
UABAF5
UABAF7
UABAF6
UABAF3
UABAF1

Supp. Fig 6.

A. Sol-BG concentration in AF samples categorized by the respective repository. Each data point represents a single patient sample. B. Kaplan-Meier survival plot of stage 3, OVCA patients stratified into low (<30th percentile) and high sol-BG (>70$^{th}$ percentile) groups. A log-rank (Mantel-Cox) test was performed, p = 0.0236, median survival Low sol-BG = 47.2 months, High sol-BG = 22.0 months. C. i) Autoradiograph of patient ascites fluid treated with chondroitinase-ABC (0.4U) and/or heparinase-III (50ng/mL) followed by [$^{125}$ I]-TGF-β1 binding and crosslinking and immunoprecipitation using anti-BG antibody. ii) Quantification of GAG chain digestion in (i) plotted as % CS to %HS. D. BG ELISA of acellular ascites fluid used for experiments in Figure 6E, plotted by the concentration of sol-BG in pg/mL, de-identified with an arbitrary number representing each patient sample. E. BG ELISA of acellular ascites fluid of AF samples used for experiments in Figure 6F, plotted by the concentration of sBG in pg/mL, identified with an arbitrary number representing each patient sample.

Supplementary Table 2

### Ascites fluid sample list

| AF ID | Depository | Histology | Stage | Survival (Months) | sol-BG Concentration (pg/mL) |
|---|---|---|---|---|---|
| DAF1 | DUKE | Endometroid | 2B | NA | 1192.1 |
| DAF2 | DUKE | Endometroid | 3 | NA | 875.2 |
| DAF3 | DUKE | mucinous | 3C | NA | 378.2 |
| DAF4 | DUKE | mucinous | 3C | NA | 1437.1 |
| DAF5 | DUKE | mucinous | | NA | 1134.1 |
| DAF6 | DUKE | mucinous | 1A | NA | 810.2 |
| DAF7 | DUKE | serous | 3C | NA | 1523.1 |
| DAF8 | DUKE | serous | 3 | NA | 1509.6 |
| DAF9 | DUKE | serous | 4 | NA | 1543.6 |
| DAF10 | DUKE | serous | 4 | NA | 1281.7 |
| DAF11 | DUKE | serous | 3C | NA | 1027.8 |
| DAF12 | DUKE | serous | 4 | NA | 915.9 |
| DAF13 | DUKE | serous | 4 | NA | 1691.5 |
| DAF14 | DUKE | serous | | NA | 551.0 |
| DAF15 | DUKE | serous | 3B | NA | 657.0 |
| DAF16 | DUKE | serous | 3C | NA | 943.0 |
| DAF17 | DUKE | serous | 4 | NA | 140.9 |
| DAF18 | DUKE | serous | 1C | NA | 696.0 |
| DAF19 | DUKE | serous | 3 | NA | 595.5 |
| DAF20 | DUKE | serous | 3 | NA | 910.7 |
| DAF21 | DUKE | serous | 3 | NA | 317.2 |
| DAF22 | DUKE | serous | 3 | NA | 505.2 |
| DAF23 | DUKE | serous | 3C | NA | 1255.6 |
| DAF24 | DUKE | serous | 3C | NA | 927.0 |
| DAF25 | DUKE | NA | NA | NA | 1872.5 |
| DAF26 | DUKE | NA | NA | NA | 482.1 |
| DAF27 | DUKE | NA | NA | NA | 847.3 |
| PAF1 | Penn. State | serous | 3C | NA | 1128.5 |
| PAF2 | Penn. State | granulosa | 1A | NA | 1057.2 |
| PAF3 | Penn. State | serous | 4 | NA | 1084.0 |
| PAF4 | Penn. State | serous | 4 | NA | 1067.5 |
| PAF5 | Penn. State | serous | 2C | NA | 1064.4 |
| PAF6 | Penn. State | serous | 4 | NA | 1096.4 |
| PAF7 | Penn. State | NA | NA | NA | 982.5 |

| AF ID | Depository | Histology | Stage | Survival (Months) | sol-BG Concentration (pg/mL) |
|---|---|---|---|---|---|
| UAF1 | UAB | papillary serous | 3C | 47 | 95.7 |
| UAF2 | UAB | serous adenocarcinoma | 3C | 88 | 174.5 |
| UAF3 | UAB | HGS | 3C | 32 | 210.0 |
| UAF4 | UAB | serous | 3C | 24 | 253.8 |
| UAF5 | UAB | papillary serous | 3C | 83 | 255.3 |
| UAF6 | UAB | Pap serous | 3C | 54 | 316.4 |
| UAF7 | UAB | papillary serous | 3C | 12 | 449.0 |
| UAF8 | UAB | NA | 3C | NA | 766.3 |
| UAF9 | UAB | HGS | 3C | 11 | 770.9 |
| UAF10 | UAB | Carcinosarcom | 4 | 33 | 847.6 |
| UAF11 | UAB | HGS | 3C | 23 | 866.3 |
| UAF12 | UAB | HGS | 3C | 3 | 903.5 |
| UAF13 | UAB | Adeno | 3C | 24 | 905.8 |
| UAF14 | UAB | serous | 3C | 51 | 920.1 |
| UAF15 | UAB | serous papilary | 3C | 24 | 1003.7 |
| UAF16 | UAB | Pap serous | 3C | 10 | 1006.6 |
| UAF17 | UAB | Prim Perit | 4 | 48 | 1063.7 |
| UAF18 | UAB | HGS | 3C | 53 | 1130.2 |
| UAF19 | UAB | ovary | 4 | 7 | 1131.5 |
| UAF20 | UAB | Prim Perit | 3C | 63 | 1162.2 |
| UAF21 | UAB | Prim Perit | 3C | 50 | 1164.0 |
| UAF22 | UAB | HGS | 3C | NA | 1569.8 |
| UAF23 | UAB | serous | 3C | 13 | 1579.9 |
| UAF24 | UAB | Endometroid | 4 | 13 | 1713.9 |
| UAF25 | UAB | Pap serous | 3C | 22 | 1791.1 |
| UAF26 | UAB | Pap serous | 3C | 22 | 2632.7 |

Supplementary Table 2.

Patient ascites fluid sample list containing histology, pathology, survival, and sol-BG concentration values.
